# nucGEMs probe the biophysical properties of the nucleoplasm

**DOI:** 10.1101/2021.11.18.469159

**Authors:** Tong Shu, Tamás Szórádi, Gururaj R. Kidiyoor, Ying Xie, Nora L. Herzog, Andrew Bazley, Martina Bonucci, Sarah Keegan, Shivanjali Saxena, Farida Ettefa, Gregory Brittingham, Joël Lemiere, David Fenyö, Fred Chang, Morgan Delarue, Liam J. Holt

## Abstract

The cell interior is highly crowded and far from thermodynamic equilibrium. This environment can dramatically impact molecular motion and assembly, and therefore influence subcellular organization and biochemical reaction rates. These effects depend strongly on length-scale, with the least information available at the important mesoscale (10-100 nanometers), which corresponds to the size of crucial regulatory molecules such as RNA polymerase II. It has been challenging to study the mesoscale physical properties of the nucleoplasm because previous methods were labor-intensive and perturbative. Here, we report nuclear Genetically Encoded Multimeric nanoparticles (nucGEMs). Introduction of a single gene leads to continuous production and assembly of protein-based bright fluorescent nanoparticles of 40 nm diameter. We implemented nucGEMs in budding and fission yeast and in mammalian cell lines. We found differences in particle motility between the nucleus and the cytosol at the mesoscale, that mitotic chromosome condensation ejects nucGEMs from the nucleus, and that nucGEMs are excluded from heterochromatin and the nucleolus. nucGEMs enable hundreds of nuclear rheology experiments per hour, and allow evolutionary comparison of the physical properties of the cytosol and nucleoplasm.

## Main

The cell interior is a highly complex crowded environment that contains polymer meshes and dense colloidal solutes of a wide range of sizes(Kate Luby-Phelps 2013; Zidovska 2020). Molecular motors and dynamic polymers create an active system that is far from equilibrium. This environment strongly influences biological reactions. One example of how the cell interior impacts biochemistry is through molecular crowding effects(Zhou, Rivas, and Minton 2008). High concentrations of crowding agents entropically favor intermolecular associations, thereby accelerating reaction rates(Rivas and Minton 2018). On the other hand, excessive crowding can also dramatically decrease molecular motion. Active processes are thought to increase the effective temperature in the cell, helping to fluidize this extreme environment. Indeed, depletion of ATP can lead to glass transitions(Parry et al. 2014). However, these glassy transitions strongly depend on length-scale: molecules with sizes equivalent to or larger than the dominant crowding agent will be more affected than small particles that can move through the gaps between larger jammed particles. For instance, in the absence of ATP, the bacterial cytosol is liquid at the nanometer length-scale of individual proteins, but becomes glassy for particles at the mesoscale (tens to hundreds of nanometers)(Parry et al. 2014). Crowding was recently demonstrated to be actively regulated at the mesoscale in the cytosol due to changes in ribosome concentration, and these changes in crowding can tune large-scale molecular assembly by phase separation(Delarue et al. 2018). However, there is still limited information about mesoscale molecular crowding in other organelles, including the nucleus.

Physical characterization within the nucleus of a living cell is challenging. Tracking of synthetic chromosomal loci(Marshall et al. 1997)(Heun et al. 2001), beads larger than 100 nm(de Vries et al. 2007; Tseng et al. 2004)(Hameed, Rao, and Shivashankar 2012), and inhomogeneities in chromatin staining(Zidovska, Weitz, and Mitchison 2013) have provided rich information about the dynamics of chromatin, but there is limited information about the properties of the fluid phase of the nucleus, the nucleoplasm. One technique that can provide extensive information about soft condensed matter is microrheology, which infers the properties of materials from the motion of tracer particles. These probes should be as passive as possible to avoid difficulties in interpretation due to binding to structures within the cell. Previous approaches to microrheology relied on the introduction of non-biological probes by microinjection(K. Luby-Phelps, Taylor, and Lanni 1986; Crick, FHC and Hughes, AFW 1950) or pinocytosis(Etoc et al. 2018), but these approaches are prohibitively labor intensive, and impossible for organisms with a cell wall, (e.g. fungi, bacteria). Fluorescence recovery after photobleaching (FRAP) and fluorescence correlation spectroscopy (FCS) experiments have provided valuable information about the nanoscale properties of the nucleoplasm(Phair and Misteli 2000), but individual fluorescent proteins are too small (∼ 3 nm in diameter) to report on the mesoscale environment.

To overcome this limitation, we recently developed genetically encoded nanoparticles based on naturally occurring homomultimeric scaffold proteins fused to fluorescent proteins(Delarue et al. 2018). In particular, we have focused on encapsulins as scaffolds, which assemble into particles of 40 nm diameter in the cytosol. We called these mesoscale probes Genetically Encoded Multimeric nanoparticles, or GEMs. GEMs allow us to probe both local and global biophysical properties of the cell in high throughput. Here, we extended this technology to the study of the rheological properties of the nucleoplasm. By introducing a nuclear localization signal (NLS) to the encapsulin protein, we directed GEMs to assemble within the nucleus. We refer to these new probes as nucGEMs, and to disambiguate, in this report we will refer to the previously reported cytosolically localized particles as cytGEMs.

## Results

### Nanoparticle design

A schematic of the nucGEM described is shown in Fig. 1a. Our original GEM design was maintained; we used an encapsulin from *Pyrococcus furiosus* as our scaffold protein, and fused an m-Sapphire (T203I, A206K) variant of GFP(Zapata-Homer and Griesbeck 2003; Ehrig, O’Kane, and Prendergast 1995) to the C-terminus (Delarue et al. 2018). The A206K mutation prevents dimerization of the fluorophore (von Stetten et al. 2012). The encapsulin scaffold drives multimerization of the monomer into a T = 3 icosahedral structure(Akita et al. 2007). The topology of this domain places the N-terminus within the lumen of the assembled particle and the C-terminus on the outside, thus a dense cloud of m-Sapphire fluorophores faces the cellular environment. We empirically determined that m-Sapphire gave the brightest particles when imaged using a standard 488 nm laser illumination and emission filters designed for GFP (bandpass from 508 to 544 nm, ET525/36m, Chroma). Importantly, we determined that the optimal excitation of m-Sapphire was shifted such that it was best excited by 488 nm light in the context of GEMs, presumably due to altered photochemistry on the crowded surface of the nanoparticles. We also found that m-Sapphire photoactivated in the context of GEMs, which is convenient as particle intensity actually increases during the first few seconds of imaging. We modified the design of cytosolic GEMs by adding a nuclear localization signal (NLS) from SV40. We initially explored gene designs in the budding yeast *Saccharomyces cerevisiae*. We tried appending the NLS to either the C-terminus or N-terminus of the encapsulin monomer. We found that both designs resulted in localization of nucGEMs within the nucleus (Fig 1b; Supplementary Fig. 1a); however the C-terminal NLS appeared to lead to occasional strong interactions with the nuclear periphery, as revealed in time projections showing long residence times at the edges of the nucleus (Supplementary Fig. 1a), lower overall effective diffusion (Supplementary Fig. 1b), and stronger ergodicity breaking over time than N-terminally tagged nucGEMs (Supplementary Fig. 1c, see below and methods for further explanation). These strong interactions are probably due to high valency interactions of the multiple NLS peptides on the particle surface with the nuclear transport machinery. The N-terminal NLS signal on the other hand is ultimately buried inside the particle and therefore inaccessible to the nuclear transport machinery. Therefore, the monomer or subassemblies must be imported through the nuclear pore prior to assembly of nanoparticles within the nucleoplasm (Fig. 1a). As a result, the surface of nucGEMs is precisely the same as cytGEMs. Therefore, differences in interactions with the cell are not a concern. Fig. 1b compares the localization and tracks from cytGEMs and nucGEMs. Comparison with the localization of an mCherry-tagged Nup49 nuclear pore marker shows that nucGEMs are confined within the nucleus (Fig. 1b, right). Therefore, we settled on the N-terminal NLS as our design for nucGEMs and now have a genetically encoded tool to study the mesoscale microrheology of the nucleoplasm.

**Fig. 1:**
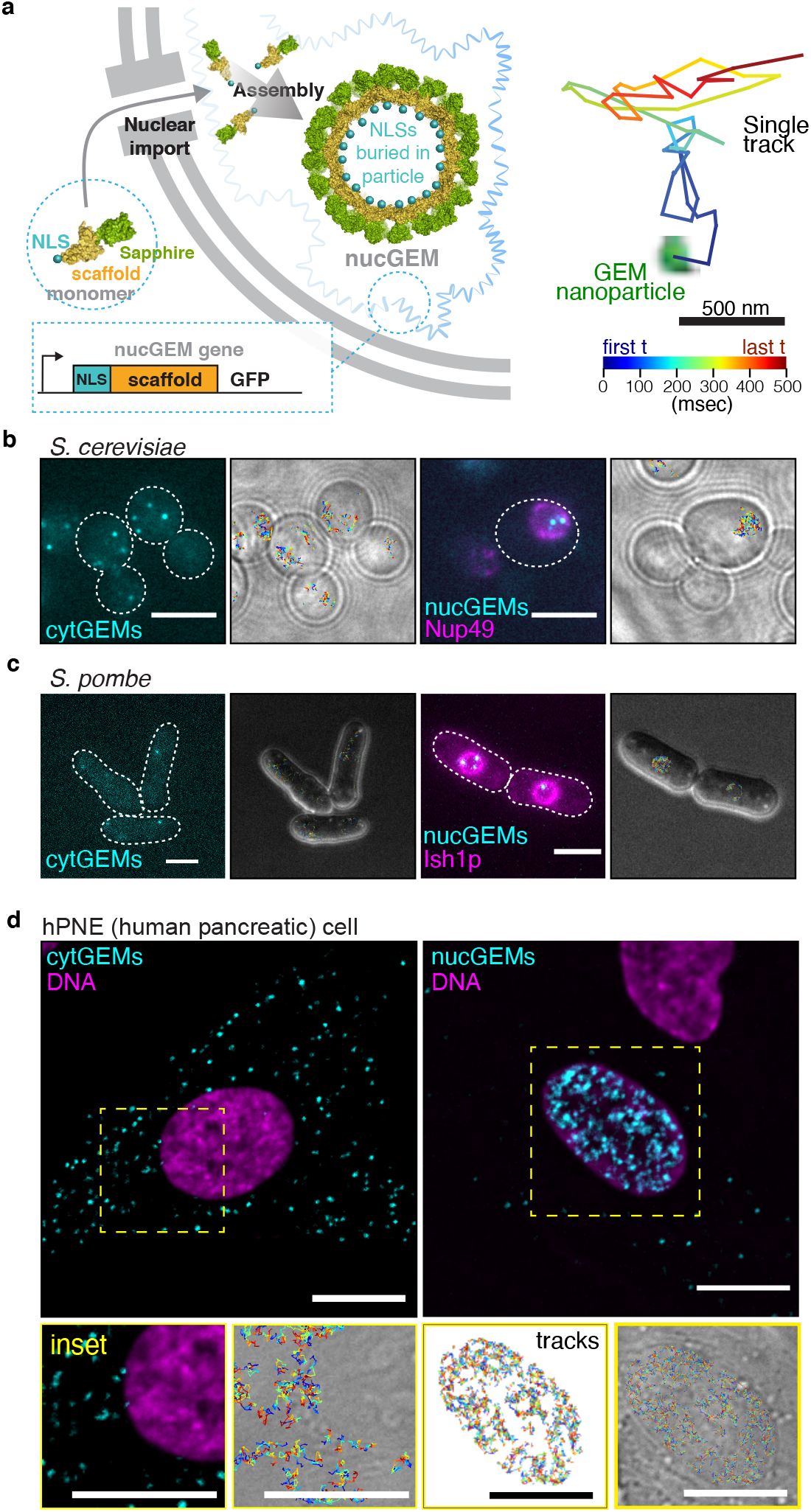
Genetically encoded nanoparticles can be targeted to assemble in the nucleus. **a**, Schematic of nucGEMs. The nucGEM gene, integrated into the genome, encodes an N-terminal nuclear localization signal on a *Pyrococcus furiosus* encapsulin scaffold, and a C-terminal m-Sapphire fluorophore. The nucGEM monomer is imported into the nucleus, and then assembles to form a 40 nm diameter nanoparticle. **b**, Representative images of cytosolic cytGEMs (cyan, left) and nuclear nucGEMs (cyan, right, with Nup49-ymRuby in magenta marking the nuclear envelope) in *Saccharomyces cerevisiae*. Also shown, tracks from movies (Supplementary Video 1) projected onto brightfield images. Scale bar represents 5 μm. **c**, Representative images of cytosolic cytGEMs (cyan, left) and nuclear nucGEMs (cyan, right, with Nup49-ymRuby in magenta marking the nuclear envelope) in *Schizosaccharomyces pombe*. Also shown, tracks from movies projected onto brightfield images. Scale bar represents 5 μm. **d**, Representative images of cytosolic cytGEMs (cyan, left) and nuclear nucGEMs (right) in human pancreatic nestin expressing (hPNE) cells. SiR-DNA dye indicates the position of the nucleus in both images. Insets show tracks from movies (Supplementary Video 2). Scale bar represents 10 μm.

### nucGEMs in fission yeast and mammalian cells

Cytosolic GEMs have been a powerful tool to compare the mesoscale physical properties of different organisms(McLaughlin et al. 2019; Delarue et al. 2018; Molines et al. 2020). Therefore, we next sought to implement nucGEMs in the fission yeast *Schizosaccharomyces pombe* and in mammalian cells. By comparison to the localization of an mCherry tagged Ish1 nuclear envelope marker, we found that 40nm-nucGEMs were also located within the nucleus of *S. pombe* (Supplementary Fig. 2). Next, we introduced 40nm-GEMs into two human cell lines, a karyotypically normal immortalized human pancreatic nestin-expressing cell line (hPNE)(K. M. Lee et al. 2003)(Fig. 1d) and the widely used HeLa epidermoid carcinoma cell line(Scherer, Syverton, and Gey 1953)(Supplementary Fig 3a). We compared the growth-rate of HeLa cells to cells stably transfected with nucGEMs and found no significant difference, indicating that the presence of these nanoparticles is not toxic (Supplementary Fig 4a). We also assessed the overall metabolic rate using PrestoBlue Cell Viability reagent and found no significant difference between control HeLa cells and nucGEM expressing HeLa cells (Supplementary Fig 4b). Previous studies found that cytGEMs are well tolerated(Carlini et al. 2020); these results indicate that nucGEMs also do not greatly perturb cell physiology.

Using the vital stain SiR-DNA (a far-red derivative of Hoechst dye, Spirochrome) we found that nucGEMs were in the nucleus of both mammalian cell lines (Fig. 1d; Supplementary Fig. 3a). However, there were also particles in the cytosol. nucGEMs are far too large to pass through nuclear pores, which have a passive diffusion size limit of around 5 nm(Mohr et al. 2009). Therefore, we hypothesized that nucGEMs might be released from the nucleus when the nuclear envelope breaks down during mitosis.

### nucGEMs are ejected from condensing mitotic chromatin

Previous work showed that cytGEMs and ribosomes are excluded from condensed prometaphase and anaphase chromatin, and incorporation into the nucleoplasm upon nuclear reassembly is further inhibited by clustering of chromosomes through a mechanism involving Ki-67(Cuylen-Haering et al. 2020). This raised the question of what happens to nucGEMs upon mitotic entry. We envisaged three possibilities: They could be trapped within condensed chromatin; excluded from chromosomes during chromatin compaction; or a mixture of these two fates. To address this question, we performed time-lapse imaging to visualize the localization of nucGEMs throughout mitosis. hPNE cells expressing nucGEM cells were synchronized in G2 with the reversible CDK1 inhibitor r3306. We imaged z-stacks of cells expressing nucGEMs motion every 15 minutes during cell division (Supplementary Video 3). Still images from a representative cell are shown in Fig. 2a.

**Fig. 2:**
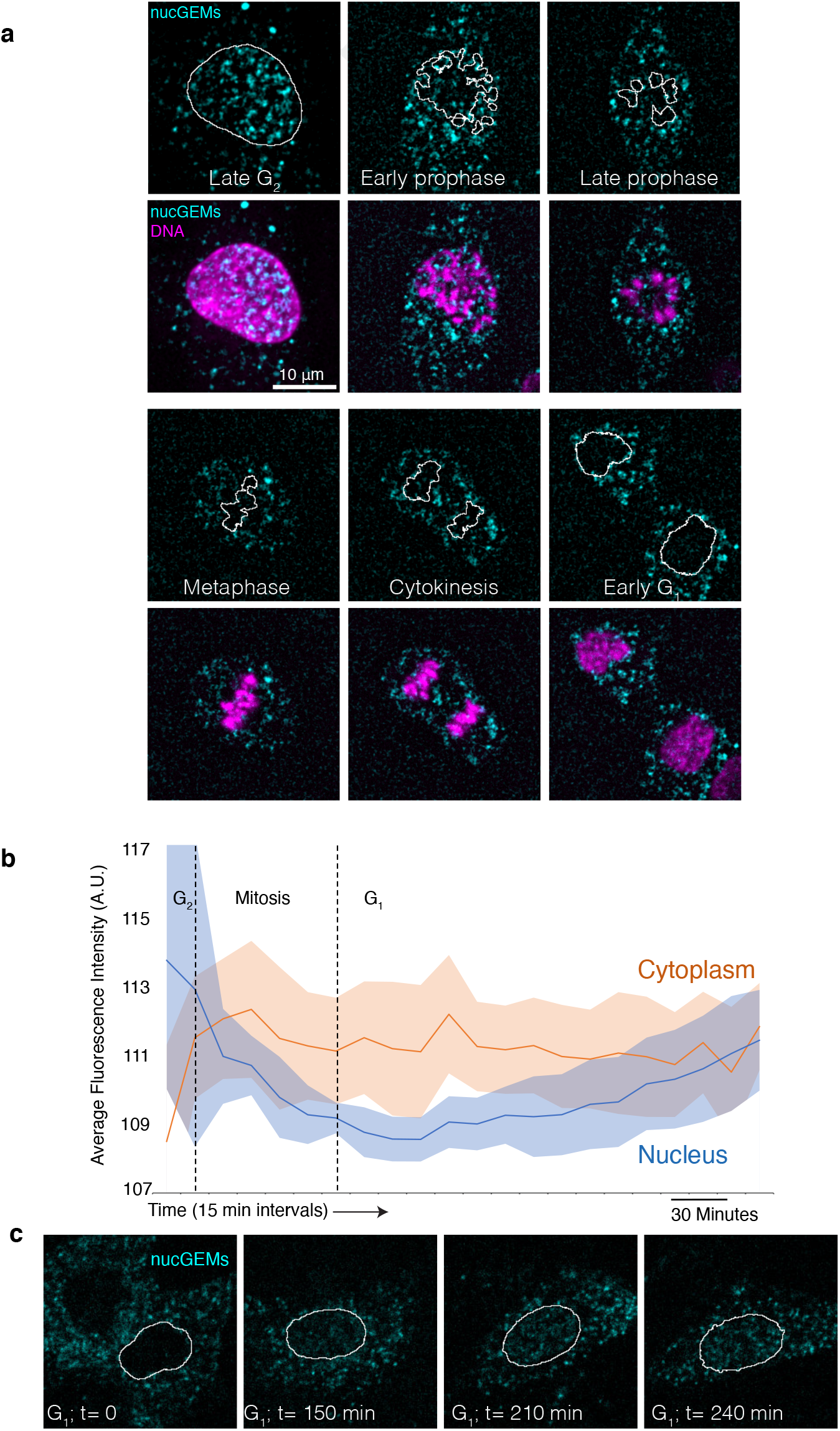
Mammalian nucGEMs assemble in the nucleus in interphase and are ejected during mitosis. **a**. Representative confocal micrographs of an hPNE cell expressing nucGEMs undergoing mitosis. nucGEMs are shown in cyan and DNA in magenta. **b**. Average fluorescence intensity of nucGEMs showing the loss and recovery of nucGEMs from the nucleus during and after mitosis. Lines represent median intensity, shaded area indicates standard deviation, n = 7. **c**, Representative confocal micrographs of an hPNE cell accumulating nucGEMs in the nucleus during the first four hours post-mitosis.

In late G2, the majority of nucGEMs are nuclear, as quantified by the average total fluorescence intensity of GEMs in the nucleus and cytosol in Fig 2b (n = 7). Upon chromosome condensation during prophase, nucGEMs became entirely excluded from chromatin and were released into the cytosol upon dissolution of the nuclear envelope. This chromatin exclusion continued throughout mitosis and, upon nuclear reassembly at telophase, very few GEMs remained in the daughter nuclei, but were instead mostly in the cytosol. Subsequently, new nucGEMs slowly assembled and accumulated in the nucleus, while the concentration of nucGEMs in the cytosol slowly decreased, perhaps due to degradation and autophagy (Fig. 2c). Together, these observations support the hypothesis that nucGEMs assemble in the nucleus, are excluded from condensing mitotic chromatin, and are ejected from the nucleus during mitosis.

The presence of nucGEMs in the cytosol and nucleus is not a problem as the nucleus and cytoplasm can be readily segmented using vital dyes or fluorescent protein markers. In fact, the presence of GEMs in both the nucleus and cytosol is very useful in comparing the properties of these compartments in the same cell, as discussed below.

### nucGEMs in *S. pombe* and post-mitotic mammalian cells

The mitotic nuclear ejection hypothesis predicts that nucGEMs will be mostly nuclear in cells with a closed mitosis, or in cells that do not divide. The nuclear envelope of *S. cerevisiae* remains intact during mitosis(Boettcher and Barral 2013). As predicted, nucGEMs are almost never observed in the cytoplasm of this organism (Fig 1b). The fission yeast *S. pombe* represents an intermediate case. The majority of these cells only contain nucGEMs in the nucleus (Fig 1c). However, we occasionally found nucGEMs in the cytoplasm of *S. pombe*, which could be due to the occasional assembly of particles prior to import, or possibly due to leakage from holes in the nuclear envelope that can appear at the spindle pole bodies during anaphase(Dey et al. 2020). Finally, consistent with the mitotic ejection hypothesis in mammalian cells, we observed that all nucGEMs were completely confined to the nucleus of post-mitotic murine neurons, and were not observed in the cytosol (Supplementary Fig. 3b).

### nucGEMs are excluded from nucleoli and preferentially explore euchromatin

The expulsion of nucGEMs from mitotic chromatin is consistent with previous work that showed that ribosomes and cytGEMs are excluded from condensed chromosomes by a mechanism involving Ki-67(Cuylen-Haering et al. 2020). This suggests that condensed chromatin could be extremely effective at excluding mesoscale particles of > 25 nm diameter (the diameter of ribosomes). However, nucGEMs do assemble and move within the interphase nucleus (Fig. 1). We additionally noticed that nucGEM tracks were mainly observed in regions of the nucleus with faint SiR-DNA straining, and appeared to be excluded from both brightly stained regions and large unstained regions. SiR-DNA binds preferentially to A/T-rich DNA, and brighter staining is thought to correspond to dense heterochromatin. On the other hand, nucleoli are typically very poorly stained by SiR-DNA and appear as dark patches. Therefore, we hypothesized that nucGEMs were excluded from interphase heterochromatin and nucleoli. First, we compared nucGEM time projections to the Nop4 nucleolar protein N-terminally tagged with mCherry in *S. cerevisiae* and found complete exclusion from the nucleolus in this organism (Fig. 3a). Next, we used immunofluorescence staining to visualize the NPM1 nucleolar marker in hPNE and HeLa cells (Fig 3b; Supplementary Fig. 3a) and again found exclusion from nucleoli. We also looked at the sc35 marker for nuclear speckles, which are also thought to be condensates(Fu and Maniatis 1990). Again, nucGEMs were excluded from nuclear speckles (Supplementary Fig. 3b). Finally, we visualized heterochromatin using antibodies against histone H3 tri-methyl lysine 9 marks (H3K9me3), and euchromatin using histone H3 acetyl-lysine 27 marks (H3K27ac) and found slight anticorrelation with the former and correlation with the latter (median Pearson R - 0.027 and +0.2316 respectively), indicating that nucGEMs are relatively excluded from heterochromatin and tend to be found mostly within euchromatic DNA (Fig. 3c). Together, these results suggest that densely packed heterochromatin and phase-separated nucleoli in interphase cells have low permeability to mesoscale particles of 40 nm diameter.

**Fig. 3:**
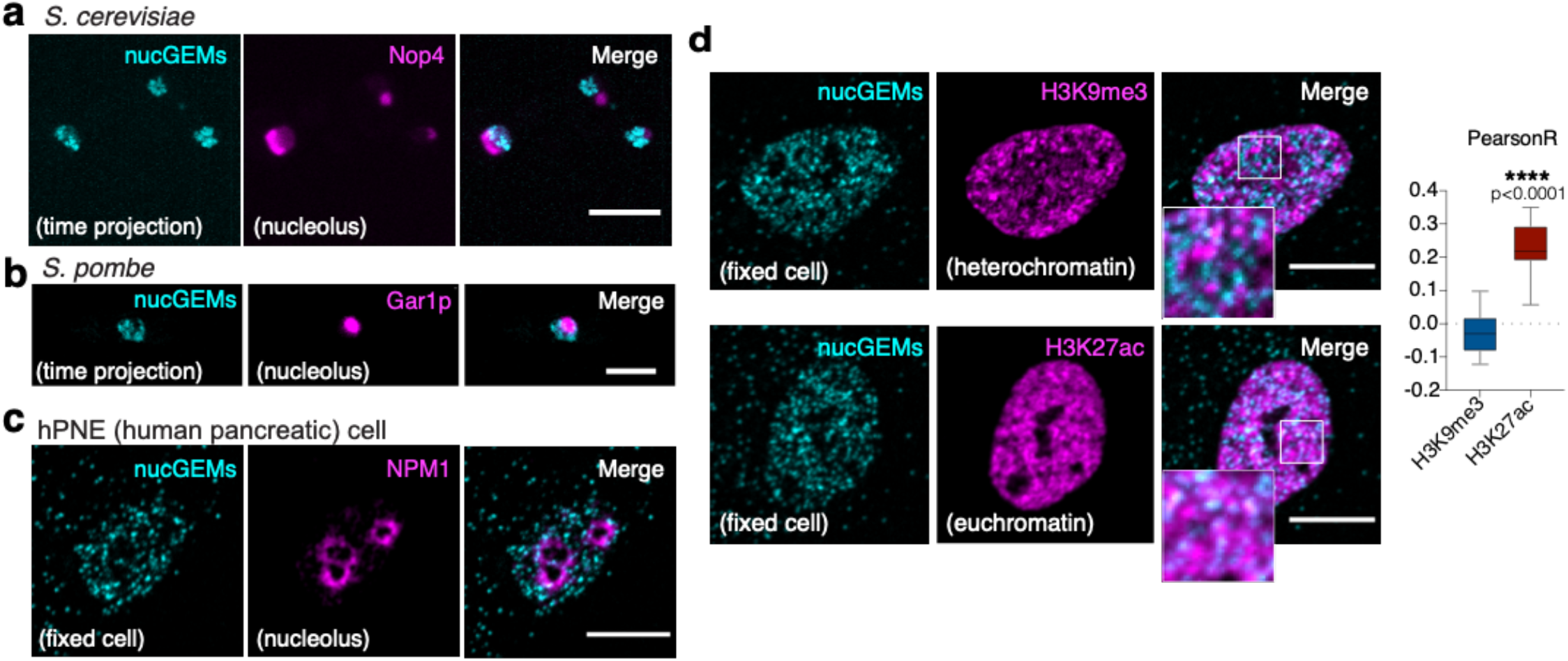
nucGEMs are excluded from nucleoli and heterochromatin. **a**. nucGEMs are excluded from the *S. cerevisiae* nucleolus. Representative images of nucGEMs (Cyan) from a time-projection of a 4 sec movie, Nop4 tagged with mCherry indicates the nucleolus (magenta), scale bar represents 5 μm. **a**. nucGEMs are excluded from the *S. pombe* nucleolus. Representative images of nucGEMs (Cyan) from a time-projection of a 4 sec movie, Gar1p tagged with mCherry indicates the nucleolus (magenta), scale bar represents 5 μm. **c**. nucGEMs are excluded from mammalian nucleoli. Representative confocal image of hPNE cell expressing nucGEMs (Cyan) stained with nucleolar marker nucleophosmin 1 (in magenta). **c**. nucGEMs are excluded from mammalian heterochromatin and enriched in euchromatin. **(left)** Representative confocal micrographs of an hPNE cell expressing nucGEMs stained with heterochromatic marker H3K9 tri-methylation or with active or euchromatin marker H3K27 acetylation (in magenta). **(right)** box plot of Pearson correlation coefficients of image pixel intensities for nucGEMS and H3K9me3 (n=18) or H3K27ac (n=17); Boxplot-whiskers represent min and max value; p-value from a Student’s t-test (**** p< 0.0001). Scale bars represent 10 μm.

### nucGEMs probe the mesoscale properties of the nucleoplasm

The nucleus has been reported to maintain a lower mass density than the cytosol by volume scaling throughout the cell cycle, while the nucleolus has the highest mass density of any compartment(Kim and Guck 2020). In addition, the nucleus has been reported to be less crowded than the cytosol at the nanometer length-scale of single GFP molecules(Phair and Misteli 2000). However, there is very little information regarding the mesoscale properties of the nucleoplasm. We therefore collected large datasets to compare the rheological properties of the nucleoplasm and cytosol in *S. cerevisiae, S. pombe*, and the hPNE and HeLa mammalian cell lines. As described above, nucGEMs appear to be excluded from heterochromatin and nucleoli, therefore we believe they are reporting on the physical properties of the nucleoplasm. We therefore refer to the nucleoplasm and cytosol in our subsequent discussion.

Data for *S. cerevisiae* and hPNE are shown in Fig. 4, and HeLa and *S. pombe* are shown in Supplementary Figs. 7 and 8 respectively. We imaged nucGEMs and cytGEMs at 100 Hz (100 frames per second) using a spinning-disk confocal for mammalian cells and Highly Inclined Thin Illumination microscopy (HILO, also referred to as incline TIRF)(Tokunaga, Imamoto, and Sakata-Sogawa 2008) for *S. cerevisiae* and *S. pombe*. We limited our analysis to particles that were tracked for longer than 10 time points and curtailed all mean-squared trajectories to this same 100 ms timescale when comparing individual tracks. Analysis of the mean square displacement (<MSD>) produced by time-averaging of these particle trajectories at a timescale of 100 ms allowed for determination of the effective diffusion constant at 100 ms (D_100ms_) of each particle(Delarue et al. 2018). We explicitly defined both effective diffusion coefficient and the anomalous exponent at the 100 ms time-scale because both of these parameters are dependent on time-scale (Supplementary Fig. 7), presumably due to the heterogeneous structure of the cytosol and nucleoplasm, which is thought to cause both local caging<u>(Chubynsky and Slater 2014)</u> and contribute active motion through metabolism(Parry et al. 2014).

**Figure 4:**
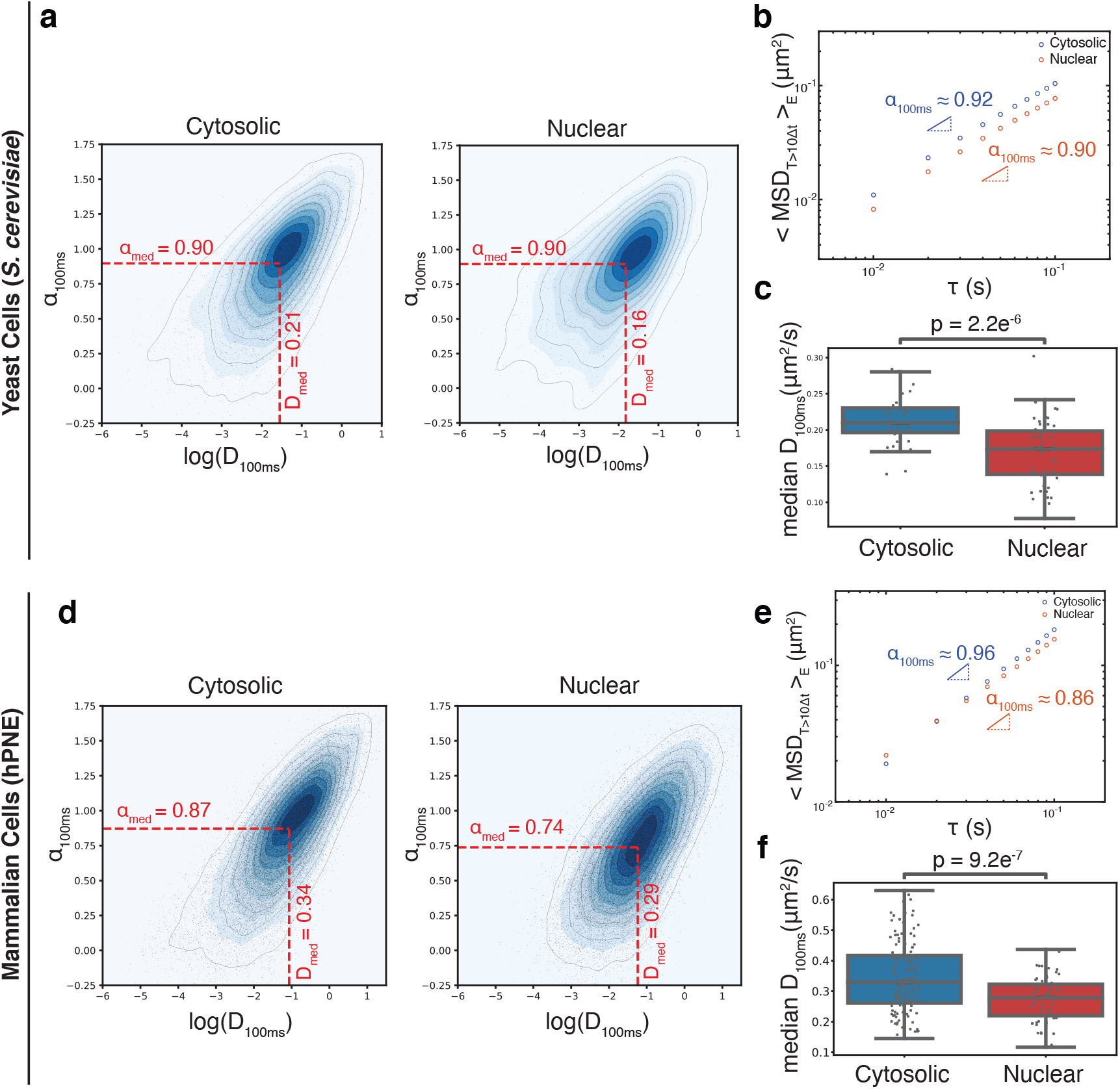
nucGEMs enable comparison of the mesoscale rheology of the nucleus and cytosol. Quantification of nucGEM movement revealed distinct mesoscale rheological properties of the nucleoplasm and cytosol in both yeast (*S. cerevisiae*, **a-e**), and mammalian cells (hPNE, **f-h**). **a**. Density plot of effective diffusion coefficients at 100 ms, versus anomalous exponents at 100 ms for individual GEM trajectories that had at least 10 time points both in the cytosol (left) and nucleoplasm (right) of *S. cerevisiae*, with their median values highlighted by red dashed lines. **b**. Ensemble- and time-averaged mean-squared displacement (MSD) versus time delay () with fits to determine values for nuclear (red) and cytosolic (blue) GEM trajectories in *S. cerevisiae* that had more than 10 time points. In **a-b**, n=10,706 (cytosolic) and n=2,969 (nuclear) trajectories. **c**. Box plots of the median effective diffusion constants (D_100ms_) of trajectories from single video fields of view of *S. cerevisiae* cells; n=36 (cytosol) and n=51 (nucleus). The horizontal lines in the boxes represent the 25^th^, 50^th^, and 75_^th^_ percentile values with whiskers extending to points that lie within 1.5 times the 25-75^th^ interquartile range. P- values are from a Student’s t-test to assess statistical differences between cytosolic and nucleoplasmic diffusion. **d**. Density plot of effective diffusion coefficients, versus anomalous exponents for individual GEM trajectories with more than 10 time points both in cytosol (left) and nucleoplasm (right) of human pancreatic (hPNE) cells. **e**. Ensemble- and time-averaged mean-squared displacement (MSD) versus time delay () with fits to determine values for nuclear (red) and cytosolic (blue) GEM trajectories in hPNE cells with more than 10 time points. In **d- e**, n=32,339 (cytosolic) and n=32,971 (nuclear) trajectories. **f**. Box plot of the median effective diffusion constants (D_100ms_) of trajectories from individual hPNE cells; n=127 (cytosol) and n=59 (nucleus).

For all organisms, there is significant variation in the mobility of individual particles in both the nucleus and the cytosol (Figs. 4a, d; Supplementary Figs. 8-11). Short tracks can give significant statistical noise, even for pure Brownian motion in a homogenous medium. Therefore, we compared our data to Brownian simulations of the same number of tracks (Supplementary Fig. 10) and found significantly more variation in our data, indicating true heterogeneity in the nucleoplasm and cytosol. In contrast to previous results at the nanoscale in mammalian cells(Phair and Misteli 2000), both individual trajectory analysis and ensemble-time averaging analysis show that, at the mesoscale in *S. cerevisiae* and mammalian cells, median nucGEM mobility measured at the 100 ms time-scale is lower in the nucleus compared to the cytGEMs in the cytosol (Fig. 4a, 4c, 4d and 4e). This lower mobility could indicate more frequent collisions with crowders (e.g. ribosomes in the cytosol) or cellular structures (such as chromatin in the nucleus) that impede motion at the 40 nm length-scale. Comparison of median α values from individual trajectories at the same 100 ms time-scale (α_100ms_) showed no significant difference between the anomalous exponent in the nucleus and cytosol (Fig. 4a). However, for *S. cerevisiae* (Fig. 4b) and human cell lines (hPNE, Fig. 4e, and HeLa, Supplementary Fig. 8b), ensemble-time averaging analysis for all trajectories with length greater than 10 (α_100ms_), gave a lower α_100ms_ in the nucleus. One interpretation for this lower α_100ms_ could be a higher level of confinement in the nucleus. This difference remains consistent over an order of magnitude of time-scales when analyzing minimum trajectory lengths of 20, 50, or 100 for *S. cerevisiae* (Supplementary Fig. 7a).

In contrast to *S. cerevisiae* and mammalian cell lines, the D_100ms_ of GEMs within the *S. pombe* nucleoplasm was higher than that within the cytosol (Supplementary Fig. 8a, c), although the nucleus still the cytosol (Supplementary Fig. 8b). It will be interesting to see how these displayed a smaller α_100ms_ than differences in nuclear rheology may reflect differences in nucleoplasmic composition and architecture.

Subdiffusive behavior (α_100ms_ < 1) can arise for multiple reasons including interactions with the cell and local or global caging of particles(Meroz and Sokolov 2015). To assess possible origins for the subdiffusive motion of GEMs, we analyzed angle correlations in our data compared to simulated Brownian motion. The angle correlations of GEMs in both the nucleus and cytosol were significantly different from the randomized angles, indicating deviation from Brownian motion (Supplementary Fig. 11b). At time scales greater than 20 ms, angle correlations were negative, consistent with local confinement forcing particles to reverse their direction. Interestingly, angle correlations were positive for very short time-scales, suggesting slightly ballistic behavior, consistent with external non-equilibrium forces imposed by active matter.

To investigate the degree of non-specific interactions between GEMs and subcellular structures, we investigated the ergodicity of our data, which is the difference between the effective diffusion D_eff_ obtained from time- versus ensemble- (spatially) averaged trajectories. Ergodicity breaking (EB) can be indicative of interactions with the cellular environment or local heterogeneities. This phenomenon can be quantified with an ergodicity breaking parameter, which is zero when there are no interactions or heterogeneities and becomes higher with stronger interactions or with greater spatial heterogeneity. We found that the D_eff_ from time-averaging has much broader distribution compared to D_eff_ from ensemble-averaging, which revealed breaking of ergodicity in both the nucleus and the cytosol (Supplementary Fig. 7d, 7f, 8d, 9d). The EB parameters were similar in both the nucleus and cytosol (Supplementary Fig. 7e, 7g, 8e, and 9e), which could suggest a similar degree of heterogeneity, or of non-specific interactions of GEMs with other structures, in these two compartments. *S. pombe* was the exception (Supplementary Fig. 9e), where ergodicity breaking increased more rapidly with time in the cytosol than in the nucleus. This might be due to previously reported strong heterogeneities in the cytosol, which is significantly denser at the tips than the center of cells<u>(Odermatt et al. 2021)</u>.

Together, these results suggest that the mobility of mesoscale (40 nm) GEMs is distinct in the nucleus and cytosol. The motion in both compartments is likely affected by crowding, confinement, and non-specific interactions. Importantly, the motion of mesoscale particles in mammalian cell lines is more rapid in the cytoplasm than the nucleus, but the opposite is true for nanoscale particles(Phair and Misteli 2000). This highlights the importance of investigating the physical properties of cells at multiple length-scales.

## Discussion

The mesoscale properties of the cytosol have been probed with well-defined genetically-encoded(Delarue et al. 2018) and non-biological(Etoc et al. 2018) nanoparticles, but there was previously very limited information for the nucleoplasm due to a lack of tools for mesoscale microrheology. Larger 100 nm nanospheres were previously used to discover important viscoelastic properties of the nucleus(Tseng et al. 2004), but we believe that these particles are large enough that they are reporting on chromatin properties rather than the fluid phase of the nucleoplasm. Synthetic(D. S. W. Lee, Wingreen, and Brangwynne 2021) and naturally occurring(Xiang et al. 2021) condensates have provided some of the best insights to date in mammalian cells and bacterial nucleoids respectively, but these probes do not assemble to a defined size and are derived from *Eukaryotic* proteins, leading to strong interactions with the cellular environment, as indicated by strongly subdiffusive behavior. We developed nucGEMs to surmount these limitations: they assemble to a defined size and geometry and are relatively passive, with few specific interactions beyond electrostatic interactions from the charge of the fluorescent protein. Furthermore, nucGEMs are easy to use: no microinjection or laborious sample preparation is required, allowing the high throughput characterization of many cells in diverse conditions, and enabling the rheological characterization of many species for the first time. For example, it is impossible to use microinjection or micropinocytosis to introduce particles into microorganisms that have cell walls, including powerful genetic systems such as *S. cerevisiae* and *S. pombe*, but we easily introduced nucGEMs into these organisms. Moreover, because nucGEMs move rapidly, a few seconds of imaging generates thousands of traces to characterize the mesoscale physical properties of the nucleoplasm at high throughput and sub-cellular resolution. Finally, after assembly nucGEMs have precisely the same size and surface properties as our previously reported cytosolic GEMs(Delarue et al. 2018), allowing meaningful comparison of the mesoscale rheological properties of the cytosol and nucleoplasm.

There are limitations in the physical interpretation of microrheology data for both technical and biological reasons. The main current technical limitation is that the data are two dimensional. Good tracking requires an imaging rate of 100 Hz (10 ms frame-rate). This rapid imaging means that experiments are high-throughput, but it is currently difficult to obtain three dimensional data with standard microscope configurations (although engineered point-spread functions are a promising potential solution(Pavani et al. 2009)). Therefore, our imaging was mostly limited to a two-dimensional plane, and as a consequence, track-lengths are often terminated by particles going out of focus. We restrict our analysis to tracks of greater than ten time-steps, but for meaningful comparison between tracks, we also curtail all tracks to this length. There is significant statistical (sampling) noise from these relatively short tracks, which is certain to contribute to the spread of effective diffusion coefficients at 100 ms (D_100ms_) and anomalous exponents at 100 ms (α_100ms_) presented in Figs. 4a and d.

Therefore, it is necessary to compare our results to simulations to evaluate true heterogeneity.

We caution against overinterpretation of the anomalous exponent α. We define both effective diffusion and the anomalous exponent at a single time-scale (100 ms) because both are variable with timescale (Supplementary Fig. 7). Different physical phenomena dominate at longer and shorter timescales. For instance, diffusion in a colloidal system is strongly time-dependent: at very short timescales, the effective diffusion coefficient is dictated by the solvent, but on longer timescales, decreases to a steady value set by collisions with crowders. In the transition between these two regimes, there is a mixture of behaviors, with rapid diffusion followed by collisions with crowders that locally and temporarily confine the particle, giving a subdiffusive behavior with an anomalous exponent *α* of below 1. Additionally, when diffusion occurs in a spatially confined environment, both effective diffusion coefficient and *α* progressively drop to 0. Thus, two non-mutually exclusive reasons for *α* values below 1 are local binding, and local or global confinement by steric interactions. It is impossible to design a completely passive particle that does not have some interactions with the cellular environment because of the enormous complexity and diversity of constituents of the cell. The surface properties of GEMs are largely defined by the properties of the densely arrayed fluorescent proteins that face toward the cellular environment. This presents a negatively charged surface that will necessarily undergo electrostatic interactions with positively charged structures in the cell. However, it has been reported that a negative surface charge is far more favorable than a positive charge in this respect(Schavemaker, Śmigiel, and Poolman 2017), perhaps because the most abundant cellular structures (ribosomes in the cytosol, nucleic acids in the nucleus) are also negatively charged. Moreover, small net charges of proteins are likely to be negligible compared to solvent friction, and could be modeled by an effective increase in the friction (Makarov and Hofmann 2021). Thus, the most likely explanation for the observed *α* value below 1 is an effective confinement, which could be due to local crowders or subcellular structures (e.g. confinement by local membranes), as well as the natural physical boundaries of the cell / nucleus.

Finally, there are currently limitations in physically interpreting microrheology data related to the fact that the cytosol and nucleoplasm are not homogenous materials, but rather complex, non-equilibrium environments. The cell is highly dynamic, and local rearrangements will constantly modify physical properties invalidating mean field assumptions and potentially explaining the observed ergodicity breaking. However, the large amounts of data that we can now access presents exciting possibilities for the development of new theoretical and simulation frameworks to understand the material properties of the cell and the impact that this unusual physical environment might have on molecular biology.

Using nucGEMs, we found that the nucleus is more crowded than the cytosol at the mesoscale in mammalian cells, which contrasts with the nanoscale where the converse is true(Phair and Misteli 2000). We find that nucGEMs are excluded from heterochromatin and the nucleolus, supporting the hypothesis that some complexes of similar size (e.g. RNA polymerase, mediator, BAF) may be physically excluded from this dense chromatin. In support of this idea, nucGEMs are ejected from the nucleus of mammalian cells at every mitosis, highlighting the dramatic cellular organization that can be achieved through changes in local material properties. This mitotic ejection also provides a mechanism to remove inappropriate particles or aggregates from the nucleus that are too large to pass through nuclear pores. Thus, the nucleoplasm can effectively be cleaned every cell cycle.

We now have a powerful technology that can investigate the mesoscale rheology of the nucleoplasm in high-throughput, thus enabling discovery of mechanisms that control crowding and activity, and the impact of these physical properties on biological processes. The size and surface properties of nucGEMs and cytGEMs are identical, allowing comparison of the physical properties of these compartments. Finally, cytGEMs and nucGEMs can be implemented in multiple organisms across the tree of life, including those with a cell wall, providing insights into the evolution of fundamental physical properties of the cell.

## Acknowledgements

We thank Diane Simeone for the gift of human pancreatic nestin expressing (hPNE) cells, Jef Boeke for HeLa and HEK293 cells, and Mario Pende for Neural Progenitor Cells (NPCs). We thank all members of the Holt lab for assistance and feedback. This work was funded by NIH R01 GM132447 and R37 CA240765, the American Cancer Society – Cornelia T. Bailey Foundation Research Scholar Grant, RSG-19-073-01-TBE, the Pershing Square Sohn Cancer Award, the Chan Zuckerberg Initiative, and the Air Force Office of Scientific Research (AFoSR) grant FA9550-21-1-3503 0091.

## Methods

| Reagent | Source | Identifier |
| --- | --- | --- |
| **Plasmids** |  |  |
| psPAX2 | Addgene | 12260 |
| pMD2.G | Addgene | 12259 |
| pINO4-SV40NLS-PfV-Sapphire-HO | Holt lab | pLH1459 |
| pPGK1-Cas9-tPGK1-URA3 | Tom Ellis lab | pLH1460 |
| gRNA-HO | Holt lab | pLH1461 |
| pRS305-pINO4-PfV-Sapphire | Holt lab | pLH497 |
| pFA6a-link-ymRuby2-SpHIS5 | Holt lab | pLH1662 |
| pRS305-pINO4-SV40NLS-PfV-Sapphire | Holt lab | pLH941 |
| pRS305-PINO4-PfV-Sapphire-SV40NLS | Holt lab | pLH637 |
| pUBC-SV40NLS-PfV-Sapphire-IRES-H2B-mCherry | Holt lab | pLH1559 |
| pUBC-PfV-Sapphire-IRES-H2B-mCherry | Holt lab | pLH1876 |
| leu1-32::pREp41X-SV40NLS-PfV-Sapphire | Chang lab |  |
| leu1-32::pREp41X-PfV-Sapphire | Chang lab |  |
| **Chemicals and consumables** |  |  |
| VECTASHIELD® Mounting Medium with DAPI | Vector laboratories | H-1800-10 |
| FuGENE® HD Transfection Reagent | Promega | E2312 |
| Ro-3306 | MedChemExpress | HY-12529 |

### Plasmid construction

The pLH1559 construct was cloned from a previous mammalian codon optimized plasmid (Delarue et al., 2018). pLH1876 was derived by restriction digestion of pLH1559 to remove the SV40NLS.

### Yeast strains

Culture: Strains were grown in synthetic complete media + 2% dextrose (SCD) according to standard Cold Spring Harbor Protocols at 30^0^C in a rotating incubator unless otherwise stated.

### Table: Strains

| Strain | Source | Identifier |
| --- | --- | --- |
| ***S. cerevisiae*** |  |  |
| BY4741, ura3Δ0, his3Δ0, leu2Δ0, met15Δ0 | Holt lab | LH2145 |
| BY4741, HO::pINO4-SV40NLS-PfV-Sapphire | Holt lab | LH4046 |
| BY4741, leu2Δ::pINO4::pINO4-PfV-Sapphire-LEU2 | Holt lab | LH4248 |
| BY4741, HO::pINO4-SV40NLS-PfV-Sapphire, Nup49-ymRuby2::SpHIS5 | Holt lab | LH4278 |
| BY4741, leu2Δ::pINO4-PfV-Sapphire-SV40NLS::LEU2 | Holt lab | LH2859 |
| BY4741, leu2Δ::pINO4-SV40NLS-PfV-Sapphire::LEU2 | Holt lab | LH3080 |
| BY4741, NOP4::pTEF2-mCherry-Nop4::NatR, his3Δ1, leu2Δ0, met15Δ0, ura3Δ0, lys+, can1Δ::GAL1pr-Scel::STE2pr-SpHIS5, lyp1Δ::STE3pr-LEU2 | SWAT library(Weill et al. 2018) | LH4152 |
| BY4741, NOP4::pTEF2-mCherry-Nop4::NatR, HO::pINO4-SV40NLS-PfV-Sapphire, can1Δ::GAL1pr-Scel::STE2pr-SpHIS5, lyp1Δ::STE3pr-LEU2 | Holt lab | LH4162 |
| ***S. pombe*** |  |  |
| h-, leu1-32::pREp41X-PfV-Sapphire, ade6-M216, ura4-D18, his3-D1 | Chang lab | FC3289 |
| h-, leu1-32::pREp41X-SV40NLS-PfV-Sapphire, ade6-M216, ura4-D18, his3-D1 | Chang lab | FC3322 |
| h-, ish1-mCherry:hphMX6, leu1-32::pREp41X-SV40NLS-PfV-Sapphire, ade6-M216, ura4-D18, his3-D1 | Chang lab | FC3330 |
| H-, c::adh21-gar1-mcherry:hph, ish1-mCherry:hphMX6, leu1-32::pREp41X-SV40NLS-PfV-Sapphire, ura4- ade6-M216 leu1-32 his3-D1 | Chang lab | FC3331 |

### Transformation

#### Mammalian cell culture and treatments

HeLa and HEK293T cells were a kind gift from Prof. Jef Boeke (Institute for Systems Genetics, NYU Langone), hTERT-immortalized HPNE cells were a kind gift from Prof. Diane Simeone (NYU Langone), and mouse neural progenitor cells (NPCs) isolated from E14.5 embryos were a kind gift of Dr. Mario Pende (INEM Paris, France). HeLa and HEK293T cells were grown in DMEM (Gibco, Cat. No. 11995073) supplemented with 10%FBS (Gemini bio-products, Cat. no. 100-106), 2mM L-Glutamine (Gibco, Cat. No. 25030-081) and Penstrep (Gibco, Cat. No.15140-122). hPNE cells were grown in RPMI 1640 (Gibco, Cat. No.11875085) supplemented with 10% FBS (Gemini bio-products, Cat. no. 100-106) and Penstrap (Gibco, Cat. No.15140-122). NPCs were grown using the NeuroCult TM Proliferation kit (Stem Cell California Inc., Cat. No. 05702) and media supplemented with 20ng/ml of human recombinant epithelial growth factor (EGF - STEMCELL Technologies, Cat. No. 78006). All cells were grown in a humidified incubator atmosphere at 37°C and 5% CO_2_. NPCs were differentiated by culturing in the absence of EGF in N2/B27 media (Neurobasal-A medium -Thermo Scientific, Cat. No. 10888022) supplemented with N2/B27 with Vitamin A (Thermo Scientific, Cat. No. 17502-048/17504044) and 0.4 mM ascorbic acid (Sigma-Aldrich, Cat. No. A8960) for at least 6 days before fixation.

#### Lentivirus production and cell transduction

HEK293T cells (9×10^6^ per 15 cm dish) were plated in antibiotic free DMEM (Gibco, Cat. No. 11995073) supplemented with 10%FBS (Gemini bio-products, Cat. no. 100-106), 2mM L-Glutamine (Gibco, Cat. No. 25030- 081). The next day, cells were transfected with transgene plasmid together with lentivirus packaging plasmids psPAX2 (Addgene, Cat. No. 12260) and pMD2.G (Addgene, Cat. No. 12259), using fuGENE HD™ transfection reagent following manufacturer’s protocol. 24 hours later, antibiotic free DMEM was replaced and supernatants collected at 48 and 72 h post-transfection and stored at 4°C. Virus titers were concentrated by centrifugation at 4,000 rcf for 40 minutes in an Amicon Ultra-15 30 KDa centrifugal filter (MilliporeSigma, Cat. No. UFC903024). Concentrated viral suspensions were aliquoted and stored at −80°C until later use. Lentivirus was introduced into cell lines of interest via reverse transduction with 1-10 μL of concentrated virus in fresh media, and replacing media after 24 hours. After cell lines stabilized, they were frozen in 10% DMSO (Sigma-Aldrich, Cat. no. D2650- 100) in FBS (Gemini bio-products, Cat. no. 100-106) and thawed for use in experiments as needed.

#### Cell cycle synchronization and imaging

200,000 HPNE cells stably expressing nucGEMs were plated in 6-well glass bottom dishes. 10uM CDK1 inhibitor Ro-3306 (MedChem Express, Cat. No. HY-12529) was added to each well and was incubated overnight (16-20 hours). On the day of the experiment, cells were mounted on a Nikon spinning disk confocal scanning microscope, equipped with a 63X/1.4 numerical aperture (NA) objective and incubator to maintain 37°C and 5% CO_2_. Cells were manually selected for imaging and imaged once in G_2_ before synchronized release. To release cells into mitosis, cells were washed 3 times with prewarmed PBS then supplied with fresh media without the drug. Time-lapse acquisition was performed with time intervals of 15 mins for 3-5 hours. Cells Undergoing mitosis were further processed and analysed using Fiji/imageJ (version 2.3.0). Images from a single focal plane were cropped and processed (subtract background, gaussian blur and adjust threshold) to generate representative images. Fluorescence intensities within the nucleus and cytoplasm were measured by segmenting the nuclear area with SiR-DNA fluorescence or hand-sampling within the cytoplasm and reported using mean gray values in the 488 nm channel. Values were averaged and plotted with standard deviation in Microsoft Excel (version 16.54).

#### Immunofluorescence analysis

Cells were fixed with 4% formaldehyde (15 min), permeabilized with 0.2% Triton X-100 in phosphate-buffered saline (PBS) for 15 min, blocked with 1% bovine serum albumin in PBS (blocking buffer) for 1 h, incubated with primary antibodies (diluted in blocking buffer) for overnight in humidified chamber at 4°C. The next day, cells were washed three times with PBS (10min interval) and then incubated in secondary antibodies (1: 400 in blocking solution) for 1 h in the dark at RT, followed by three PBS washes. Samples were mounted with VectaShield mounting medium containing 4′,6-diamidino-2-phenylindole (DAPI). Cells were incubated with SiR-DNA for 1 h prior to fixation, in cases where SiR-DNA was used as a DNA marker. Image acquisition was performed using Nikon spinning disk confocal scanning microscope, equipped with a 63X/1.4 numerical aperture (NA) objective. Images were processed using FIJI/ImageJ2 (version 2.3.0). All images unless mentioned otherwise are single optical sections of the images (step size 0.5μm). PearsonR method: Pearson correlation coefficients were calculated by comparing pixel intensities of each channel.

| Subcellular compartment | Marker | Source and catalog No. | Dilution |
| --- | --- | --- | --- |
| Heterochromatin | Histone H3 (tri methyl K9) | Abcam (ab8898) | 1:500 |
| Nuclear Speckle | SC-35 Nuclear Speckle Marker | Abcam (ab11826) | 1:250 |
| Nucleolus | NPM1 (Nucleophosmin) | Abcam (ab10530) | 1:500 |
| Euchromatin | Histone H3 (acetyl K27) | Abcam (ab4729) | 1:500 |
| Nuclear envelope | Lamin B1 | Abcam (ab16048) | 1:500 |
| Neural soma and dendrites | MAP2 | Proteintech (17490-1-AP) | 1:500 |
| DNA | SiR-DNA | Cytoskeleton (CY-SC007) | 1:5000 |
| Secondary Antibodies | Goat Anti-Rabbit IgG (H+L)-Alexa 568 | Invitrogen (A11036) | 1:1000 |
| Secondary Antibodies | Goat Anti-Mouse IgG (H+L)-Alexa 594 | Invitrogen (A11032) | 1:1000 |

#### Cell Growth and Viability Assay

For cell growth analysis, 20,000 cells were seeded in a 24-well plate. After 24hrs cells were trypsinized using TrypLE™ Express Enzyme (GIBCO, Cat. no. 12604039) and total number of live cells were counted based on Trypan Blue stain using Countess^®^ II FL Automated Cell Counter (Invitrogen, Cat. No. AMQAF2000). Metabolic activity was assayed as a measure of cell viability using PrestoBlue™ (Invitrogen, Cat. no. A13261). 20,000 cells were seeded per well in a 24-well plate and, 24, 48, or 72 hours later, PrestoBlue™ was added to cell media according to manufacturer’s instructions. After 1 hr incubation, the supernatant was transferred to a 96-well flat bottom plate and fluorescence was measured at excitation wavelength of 560 nm and emission at 590 nm using a microplate reader.

#### Highly inclined thin illumination (HILO) Imaging of *S. cerevisiae* GEMs

GEM particles were constitutively expressed from the very weak INO4 promoter in *S. cerevisiae* cells. Cells were imaged using a TIRF Nikon TI Eclipse microscope in highly inclined thin illumination mode (HILO) at 488 nm excitation with 100% power. The emitted fluorescent signals were transmitted through a 100x objective (<u>100x DIC, Nikon</u>, oil NA = 1.45, part number = MRD01905; <u>100x Phase, Nikon</u>, oil NA = 1.4, part number = MRD31901) and recorded with a sCMOS camera (<u>Zyla, Andor</u>, part number = ZYLA-4.2p-CL10). GFP filter set (<u>ET-EGFP (FITC/Cy2), Chroma</u>, part number = 49002) was embedded within the light path, which includes an excitation filter (Excitation wavelength/ Bandwidth (FWHM) = 470/40 nm), a dichroic mirror (long pass beamsplitter, reflecting < 495 nm and transmitting > 495 nm wavelength) and an emission filter (Emission wavelength/ Bandwidth (FWHM) = 525/50 nm). Each GEM movie was composed of images acquired every 10 ms for a total 4 s.

#### HILO Imaging of *S. pombe* GEMs

For cytoplasmic 40 nm GEMs, PfV encapsulin-mSapphire was expressed in fission yeast cells carrying the multicopy thiamine-regulated plasmid pREP41X-PfV-mSapphire. For nuclear 40 nm GEMs, NLS-PfV-mSapphire was inserted pREP41X. The expression of these constructs was under the control of the thiamine repressible nmt41 promoter<u>(Maundrell 1990)</u>. Cells were grown using a protocol that produced appropriate, reproducible expression levels of the GEMs: cells carrying these plasmids were grown from a frozen stock on EMM3S-LEU plates without thiamine for 2-3 days at 30 °C and stored at room temperature for 1-2 days to induce expression. They were then inoculated in liquid EMM3S-LEU with 0.1 μg/mL of thiamine (#T4625-25G, Sigma Aldrich) for partial repression of the nmt41 promoter and grown for one day at 30 °C to exponential phase. Cells were immobilized in lectin-treated μ-Slide VI 0.4 channel slides (#80606, Ibidi) and imaged in fields of 250×250 pixels or smaller using HILO TIRF illumination at 100 Hz for 10 s.

#### Confocal Imaging of GEMs

Micrographs were acquired on a Nikon Eclipse Ti Eclipse microscope mounted with Yokogawa CSU-X1 spinning disk unit, NIDAQ AOTF multilaser unit, and Prime 95B camera operating on Nikon NIS-Elements AR (v 5.21.03) software. We used CFI Apo 60x/N.A-1.49/.12 TIRF objective with a 470/40m excitation filter and ET525/36m emission filter (Chroma Technology Corp) in all mammalian acquisitions. Using a 488 nm laser the sapphire fluorophore was excited using 100% power and images were collected from a single focal plane at 100fps, binning 1, 512×512, and 8-bit pixel depth for 2 to 4 seconds.

### Quantification of mesoscale rheology

#### Time-averaged, ensemble-time-averaged mean-square displacement (MSD)

GEMs were initially tracked with the ImageJ Particle Tracker 2D-3D tracking algorithm from MosaicSuite<u>(Sbalzarini and Koumoutsakos 2005)</u> and trajectories were then analyzed with the GEM-Spa (GEM single particle analysis) software package that we are developing in house: https://github.com/liamholtlab/GEMspa/releases/tag/v0.11-beta

For every 2D trajectory, we calculated the time-averaged mean-square displacement (MSD) at different time intervals:

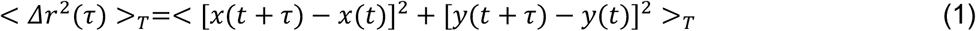

where ‘<>_*T*_’ represents time averaging for each trajectory of all displacements under time interval *τ*.

To reduce tracking error due to particles moving in and out of the focal plane, we selected particle trajectories with more than 10 time points. We then fitted the time-averaged MSD of each selected trajectory with power-law time dependence based on the first 10 time intervals (100ms). Density map of α vs. *D*_100*ms*_ can then be plotted for all trajectories (Etoc et al. 2018) (Fig. 4a, 4f and S5a, S6a).

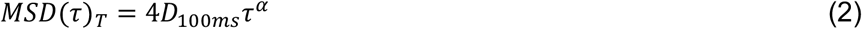

where *α* indicates diffusion property, with *α* = 1 being Brownian motion, *α* < 1 suggests sub-diffusive motion and *α* > 1 as super-diffusive motion. *D*_100*ms*_ is the diffusion coefficient with the unit of *μm*^2^/*s*^*α*^.

For better comparison of GEM diffusivity under different conditions, we also used the effective diffusion coefficient for characterization due to the unifying of its unit as *μm*^2^/*s*. Time-averaged MSD for each trajectory is fitted using a linear time dependence at first 10 time intervals:

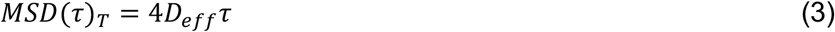

where *D*_*eff*_ is the effective diffusion coefficient for each trajectory.

We then used median value of *D*_*eff*_ among all trajectories within either each field of view for yeast cells (512×512 pixels including several yeast cells) or each individual mammalian cell and plotted as each individual dot on bar graphs for characterizing GEM mobility in different conditions (Fig. 4c, 4h, S5c and S6c).

Ensemble-time averaged MSD was also applied for better indication of *α* at each condition and was subsequently fitted with the power-law time dependence at first 10 time intervals.

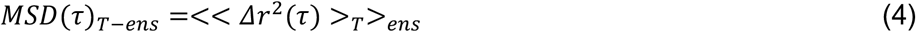

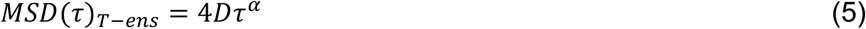

where ensemble-time averaged MSD is the ensemble-averaging among all time-averaged MSD for trajectories that are above a certain trajectory length cutoff (10 time points for most figures: Fig. 4b, 4g, S5b and S6b; 20, 50, 100 for Fig. S4a).

#### Ensemble-averaged MSD and breaking of ergodicity

At every time point, we calculated the ensemble-averaged mean-square displacement (MSD) for all trajectories based on:

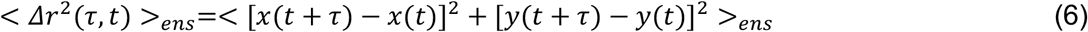

where ‘<>_*ens*_’ represents ensemble averaging for all selected trajectories with displacements under time interval *τ* at time point t.

For simplicity, we choose specific time interval *τ* = 20*ms* and directly calculate effective diffusion constant without fitting as *D*_20*ms*_ using either time-averaged or ensemble-averaged MSD at 20ms time interval (Etoc et al. 2018)(Weigel et al. 2011) (Fig. S4b, S4d, S5d and S6d).

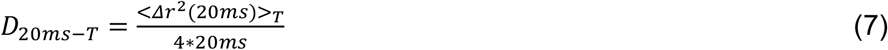

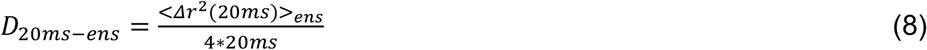

Differences in distribution of time-averaged and ensemble-averaged *D*_20*ms*_ suggests ergodicity breaking. To quantify the level of nonergodicity, we calculated ergodicity breaking parameter (EB) based on (Manzo and Garcia-Parajo 2015) (Meroz and Sokolov 2015) (Fig. S4c, S4e, S5e and S6e):

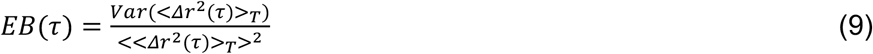

where EB value is dimensionless quantity with its numerator as the variance and its denominator as the square of mean of time-averaged MSD for all selected trajectories at time interval *τ*.

#### Angle correlation function and estimate of effective confinement size

Angle correlation function was calculated for detailed analysis of GEMs movement. For each trajectory, we calculated cosine of angle between displacements under time interval *τ*. Angle correlation function was then calculated by combining and averaging all *cos*(*θ*(*τ*)) values within each trajectory as well as among all trajectories.

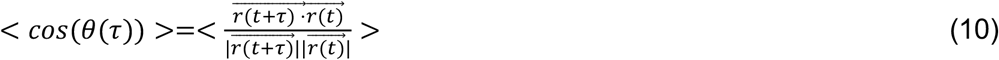

where < *cos*(*θ*(*τ*)) > = 0 suggests no angular correlation as Brownian motion, < *cos*(*θ*(*τ*)) > < 0 suggests anti-persistent angular correlation and < *cos*(*θ*(*τ*)) > > 0 indicates persistent angular correlation (Harrison et al. 2013) (Fig. 4d).

We could then calculate the characteristic time *t*_*cross*_ as the time point when < *cos*(*θ*(*τ*)) > changes from positive to negative values. *t*_*cross*_ indicated the time scale for directional GEM movements. Combining previously acquired *D*_*eff*_ for every condition, we could estimate the effective confinement size for GEM particles in both cytosol and nucleus (Fig. 4e).

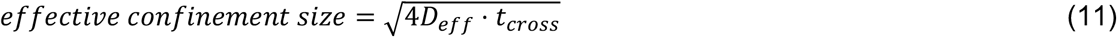

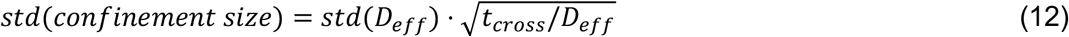

where ‘*std*’ represents the standard deviation of variables.

### Plasmid, strain and cell line availability

All plasmids will be deposited in Addgene. Yeast strains and human cell lines will be made available upon request.

## Data availability

Github: https://github.com/Shutong20/Holtlab-nucGEM-paper-data-repository

## Code availability

Matlab code: https://github.com/Shutong20/Holt-Lab-GEM-analysis

Python GEM-Spa package: https://github.com/liamholtlab/GEMspa/releases/tag/v0.11-beta

## Ethics declarations

The authors declare no competing financial interests.

