## Supplemental Figures 1 to 11 for "nucGEMs probe the biophysical properties of the nucleoplasm"

### Supplementary Information for
### nucGEMs probe the biophysical properties of the nucleoplasm

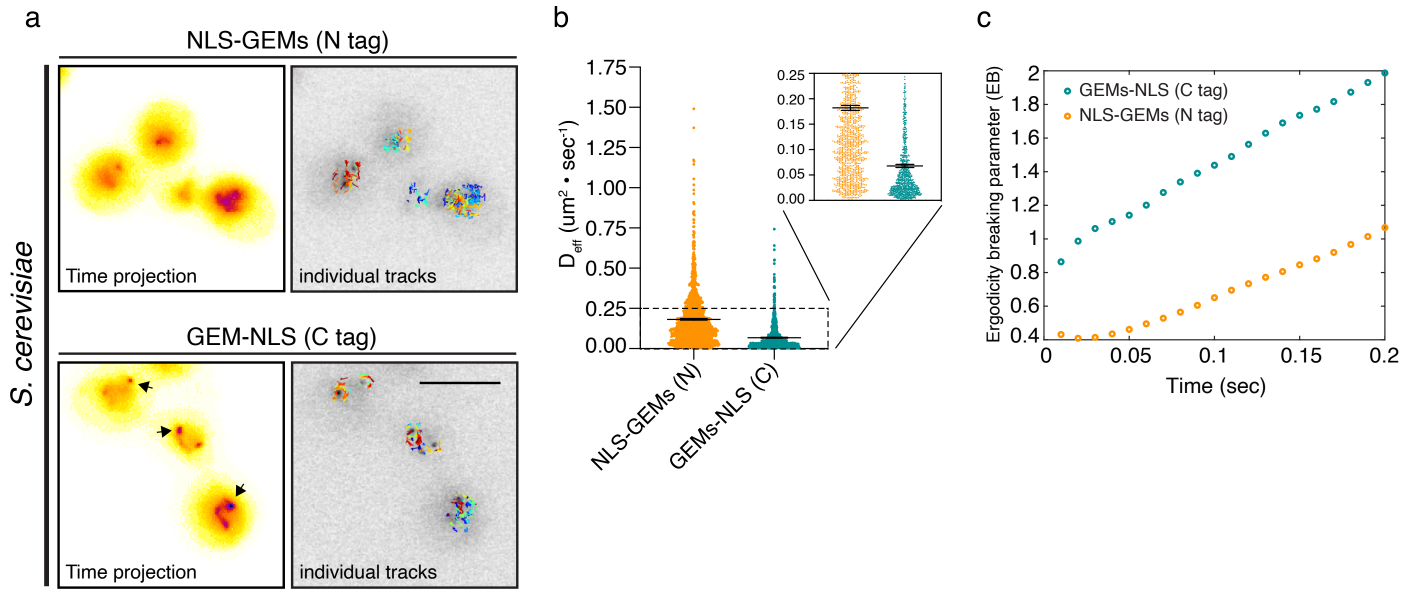


**Supplementary Figure 1: An N-terminal NLS leads to more passive nucGEMs than a C-terminal NLS.** **a.** Representative time projections from nucGEM movies where the nuclear localization signal (NLS) is tagged on the N-terminus (top panel) or C-terminus (bottom panel) of the *Pyrococcus furiosus* encapsulin protein. Arrows in the bottom panel show C-terminal NLS (nucGEM-NLS) particles that appear to transiently bind to the nuclear periphery. Individual tracks are also shown. Scale bar represents 5 μm. **b.** Effective diffusion coefficient ($D_{eff}$) calculated from movies ($n_{movies}$ = 5) of NLS-nucGEM ($n_{tracks}$ = 1245) and nucGEM-NLS ($n_{tracks}$ = 819). **c.** Ergodicity breaking parameters (EB) plotted over 200 ms for NLS-nucGEM and nucGEM-NLS suggest an N-terminal NLS reduces particle interactions.

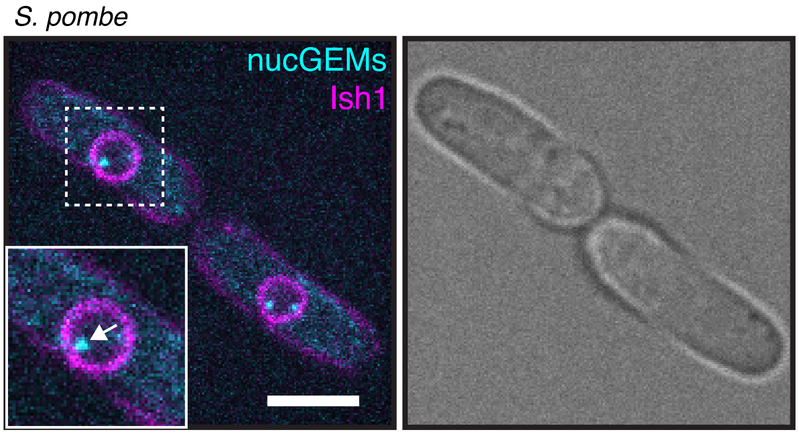


**Supplementary Figure 2: nucGEM expression in *Schizosaccharomyces pombe* a.** Representative still images of nucGEMs (cyan) in *S. pombe,* Ish1-mCherry (magenta) marks the nuclear envelope and plasma membrane. White arrow in the inset indicates nucGEM. Scale bar indicates 5 µm.

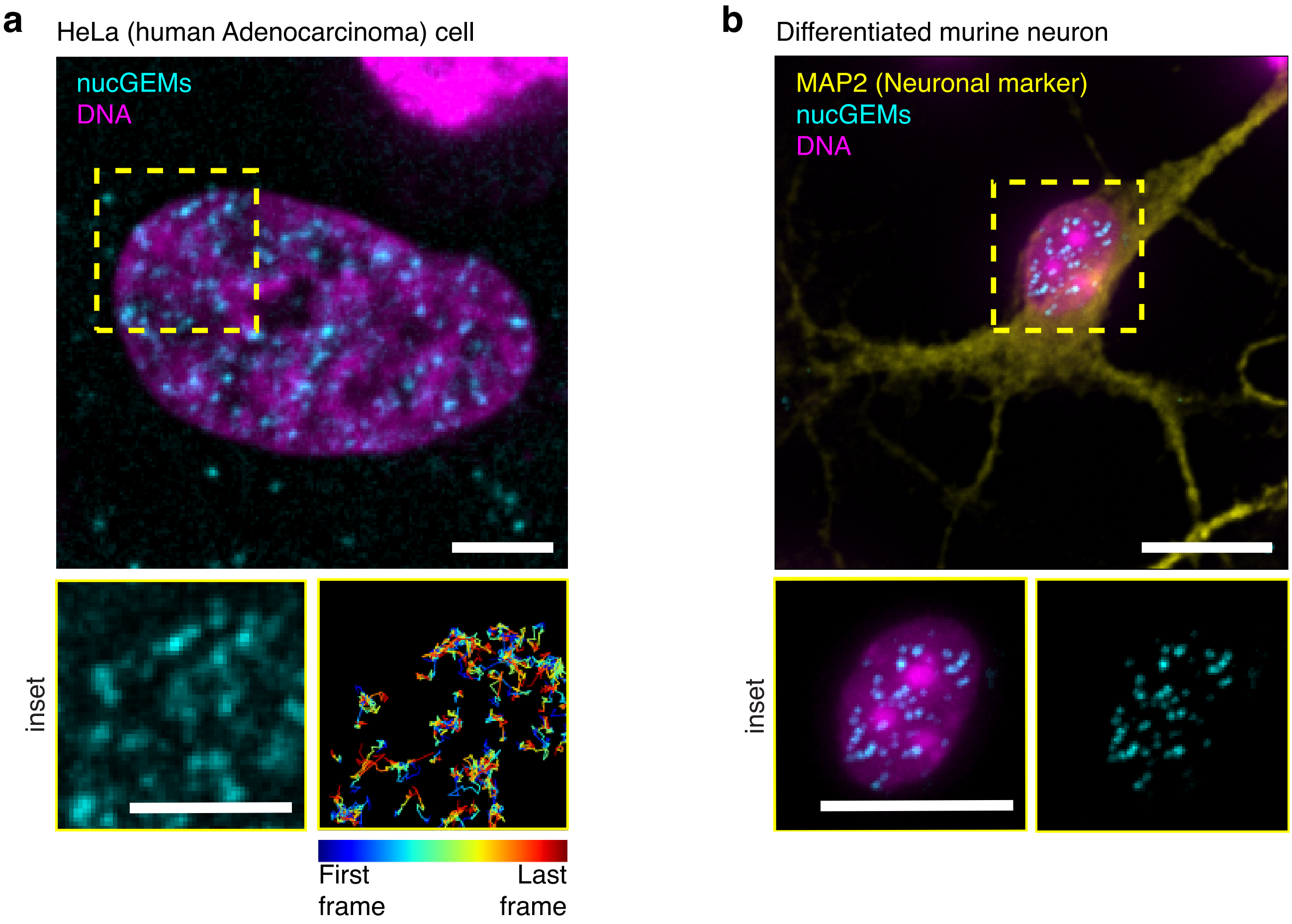


**Supplementary Figure 3: nucGEM expression in mammalian cell lines. a.** Representative image of nucGEMs (cyan) in the nucleus (SiR-DNA, magenta) of human adenocarcinoma (HeLa) cells. Inset shows temporal color coding of nucGEM diffusion represented by rainbow tracks. Scale bar represents 5 μm. **b.** Representative image of nucGEMs (cyan) in the nucleus (SiR-DNA, magenta) of a neuron differentiated from a murine neural progenitor cell, immunostained with the neuronal marker MAP2 (yellow). Inset shows higher magnification of the nucleus, and scale bar represents 10 μm.

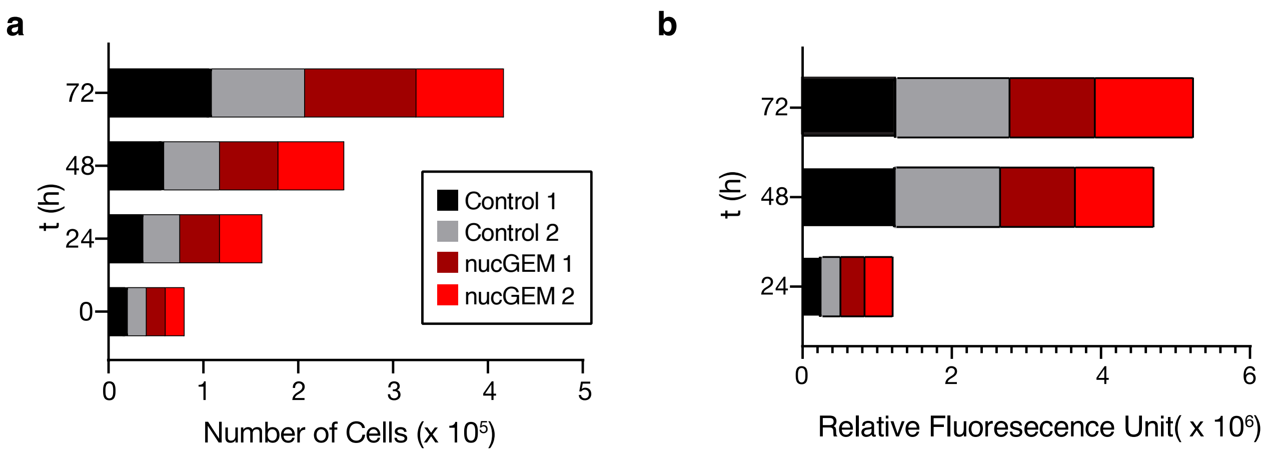


**Supplementary Figure 4: nucGEMs have no significant impact on the growth and metabolic activity of HeLa cells. a.** Growth rates of control and nucGEM-expressing HeLa cells show that nucGEMs do not impact cell growth. Equal number of cells were seeded and quantified every 24 hours using trypan blue to exclude dead cells. Two technical replicates are shown. **b.** Metabolic assay of control and nucGEM-expressing HeLa cells show nucGEMs do not impact cell viability. An equal number of cells were seeded and stained with PrestoBlue$^{TM}$ for 1 hour preceding supernatant fluorescence measurements at 590 nm.

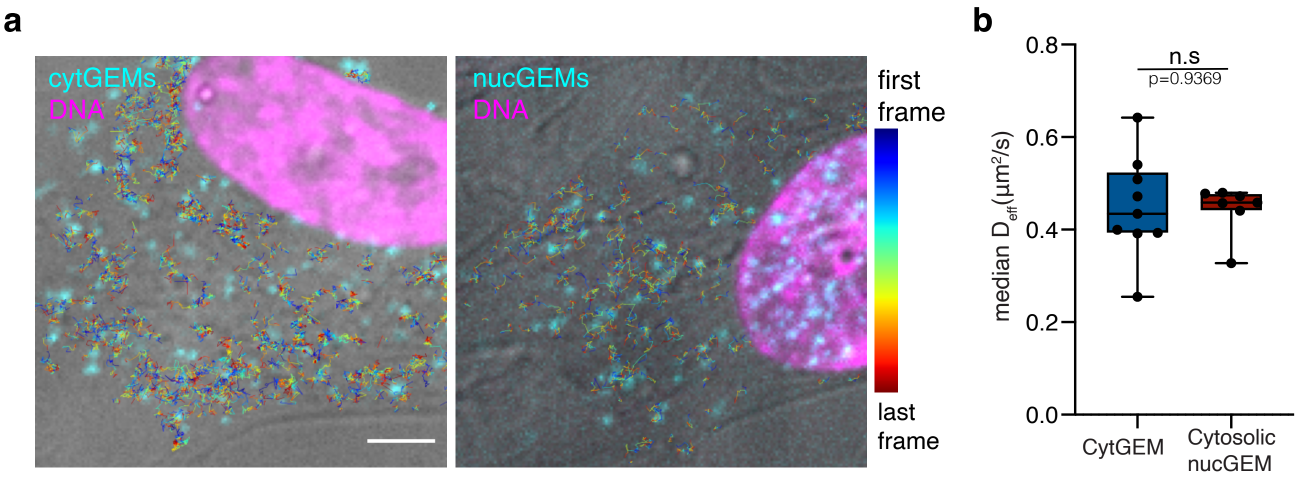


**Supplementary Figure 5: nucGEMs ejected from the nucleus behave similarly to cytGEMs. a**. Representative micrographs show hPNE cells expressing cytGEMs (left, cyan) and nucGEMs (right, cyan) with temporal color-coded tracks overlaid to indicate particle motion (SiR-DNA shown in magenta, scale bar represents 10 µm). **b**. Median effective diffusion coefficients ($D_{eff}$) of cytGEMs (blue, $n_{tracks}$ = 7663) and nucGEMs (red, $n_{tracks}$ = 3398) in individual hPNE cells ($n_{cytGEM}$ = 9, $n_{nucGEM}$ = 7) are not statistically different. Whiskers represent min and max values; *p*-values calculated using t-test.

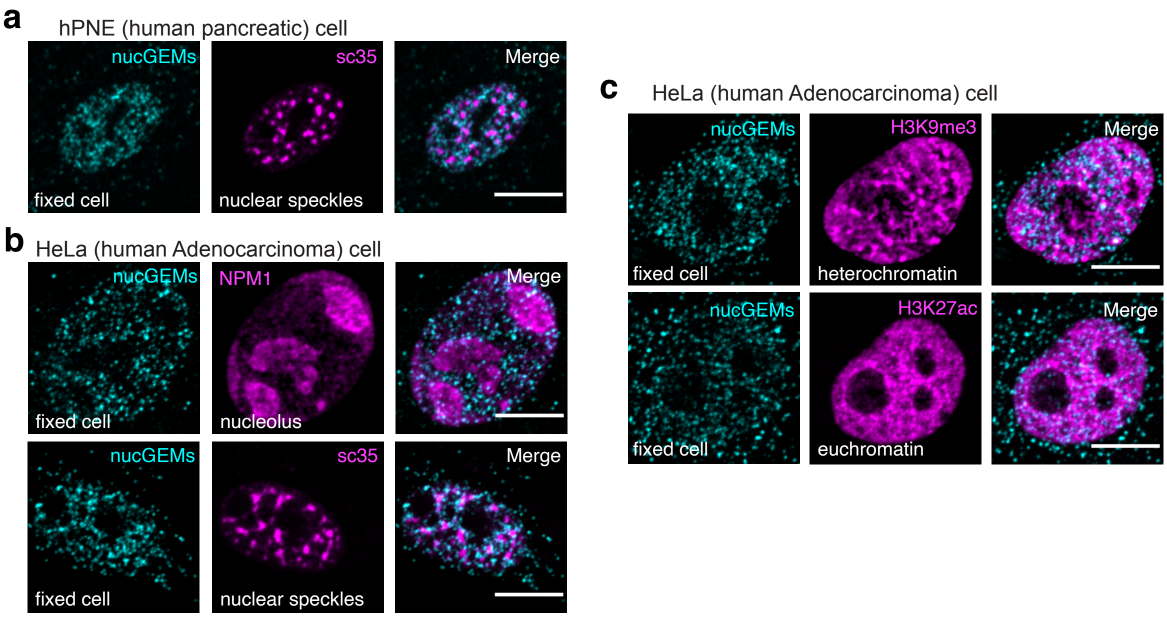


**Supplementary Figure 6: nucGEMs are excluded from nucleoli, nuclear speckles and heterochromatin in both hPNE and HeLa cells. a.** Representative confocal image of hPNE cell expressing nucGEMs (cyan) stained with nuclear speckle marker SC-35. **b.** Representative confocal images of HeLa cells expressing nucGEMs (cyan) and stained with the nucleolar marker NPM1 or with the nuclear speckle marker SC-35 (magenta, bottom). **c.** Representative confocal micrographs of fixed HeLa cells expressing nucGEMs and immunostained with antibodies that recognize the trimethylated H3K9 heterochromatic marker (magenta, top), or the acetylated H3K27 euchromatin marker (magenta, bottom). Scale bar represents 10 μm.

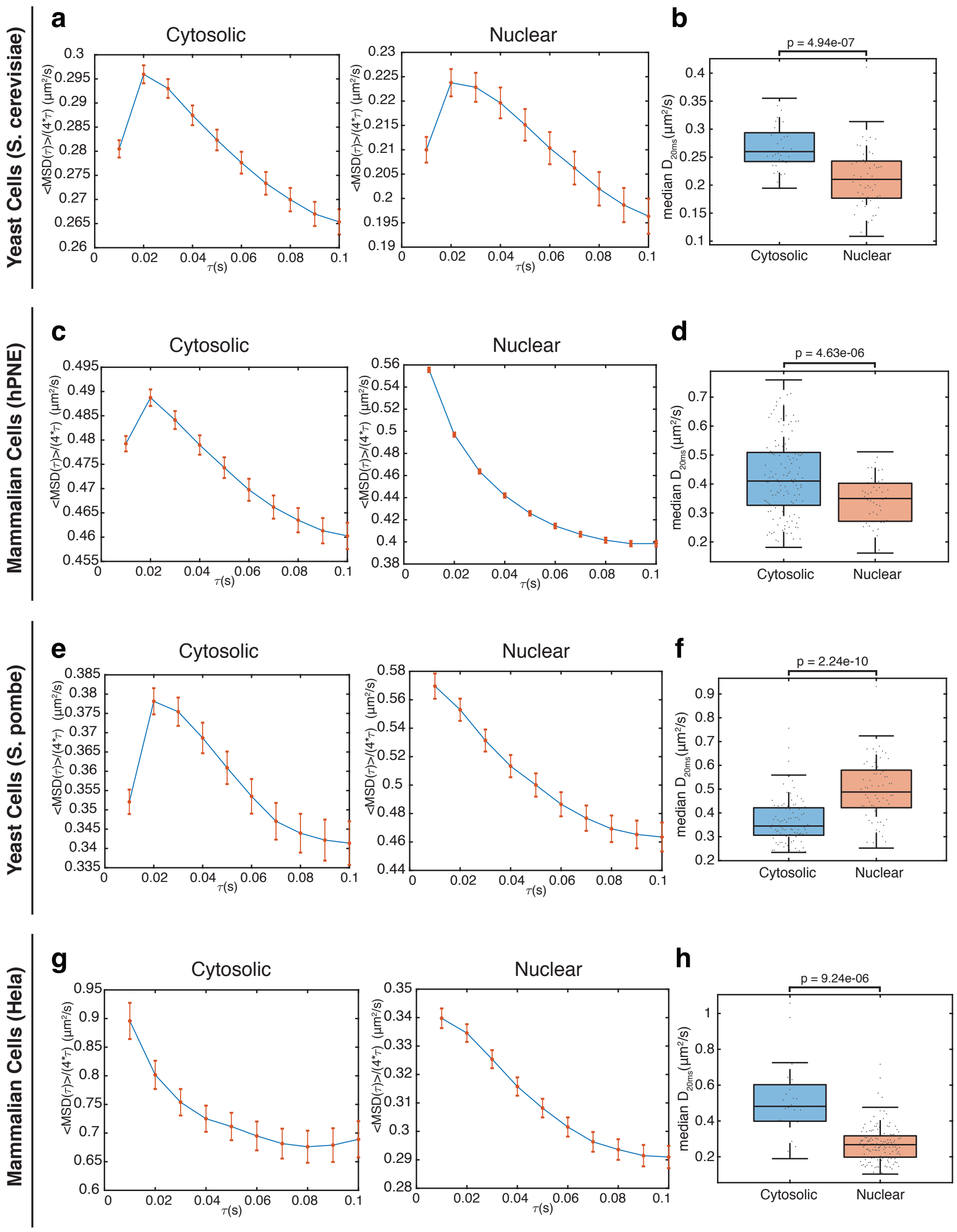


**Supplementary Figure 7: Time delay dependence of time-averaged MSD**

**Left**: Mean of time-averaged mean-square displacement over 4 time delay ($MSD(\tau)/(4*\tau)$) for all individual GEM trajectories with more than 10 time points in both the cytosol and nucleoplasm of yeast (**a:** *S. cerevisiae* $n_{cytosolic}$ = 10,706 and $n_{nuclear}$ = 2,969; **e:** *S. pombe* $n_{cytosolic}$ = 4,212 and $n_{nuclear}$ = 1,843), and mammalian cells (**b:** hPNE $n_{cytosolic}$ = 32,339 and $n_{nuclear}$ = 32,971; **g:** Hela $n_{cytosolic}$ = 839 and $n_{nuclear}$ = 8,009). Error bar represents standard error of mean. **Right**: Box plots of the median effective diffusion constants at 20ms ($D_{20ms}$) of trajectories from single video fields of view of *S. cerevisiae* cells (**b** $n_{cytosolic}$ = 36 and $n_{nuclear}$ = 51), individual *S. pombe* yeast cells (**f** $n_{cytosolic}$ = 94 and $n_{nuclear}$ = 65), individual hPNE cells (**d** $n_{cytosolic}$ = 127 and $n_{nuclear}$ = 59) and individual Hela cells (**h** $n_{cytosolic}$ = 24 and $n_{nuclear}$ = 143). P-values were derived from a Student's t-test to assess statistical differences between cytosolic and nuclear diffusion.

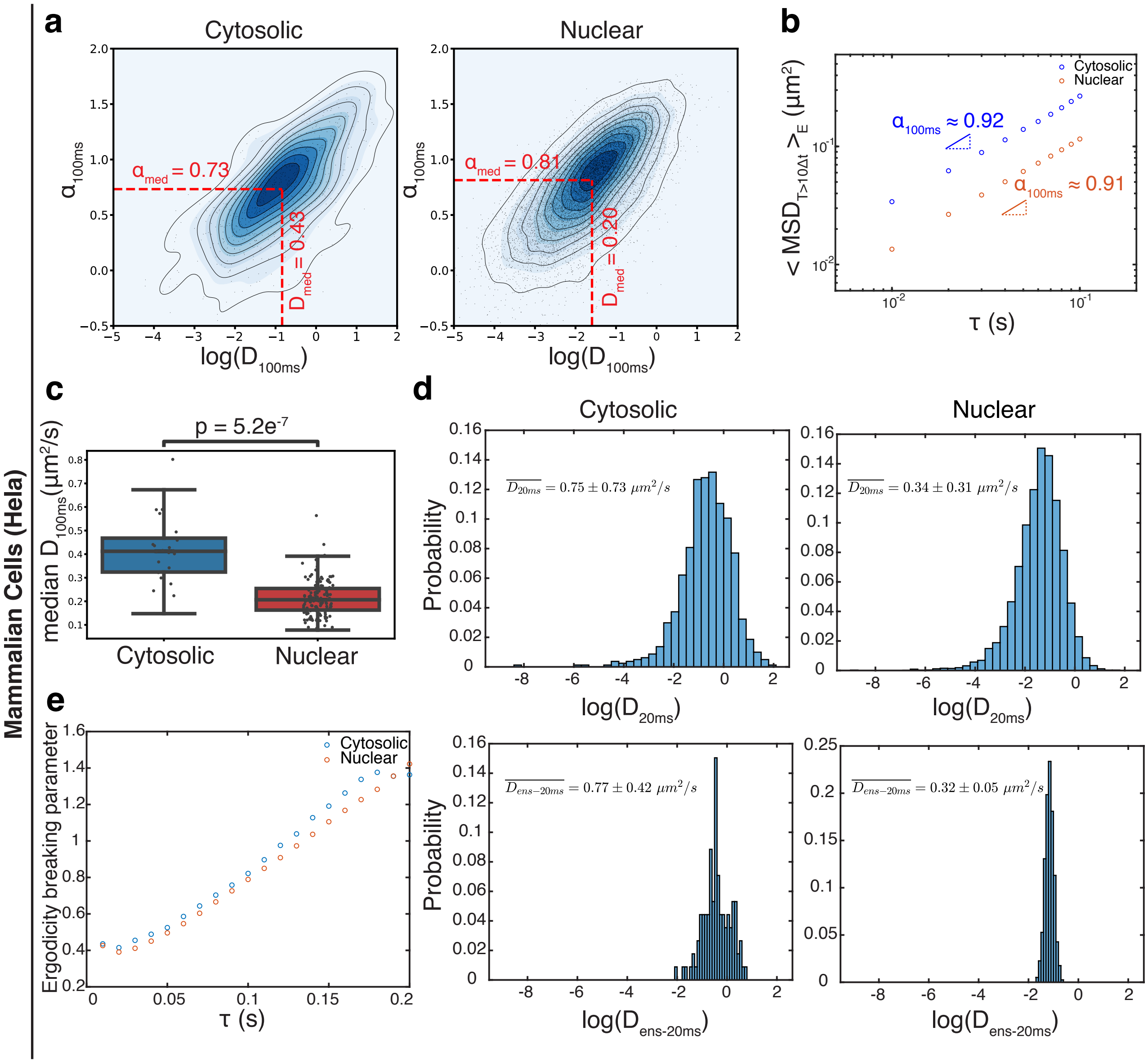


**Supplementary Figure 8: Characterization of the rheological properties of the cytosol and nucleoplasm of Hela cells. a**. Density plot of $\alpha_{100ms}$ - versus log($D_{100ms}$) for individual GEM trajectories in the cytoplasm (left) and nucleus (right) of Hela cells. **b**. Ensemble-time-averaged MSD with fitted $\alpha_{100ms}$ values for GEM trajectories in Hela cells with larger than 10 time points. In **a-b**, trajectories $n_{cytosolic}$ = 839 and $n_{nuclear}$ = 8009. **c**. Box plot of the median effective diffusion constants ($D_{100ms}$) of trajectories from individual Hela cells; $n_{cytosolic}$ =24 and $n_{nucleus}$ =143, *p*-value calculated using Student's t-test. **d**. Histograms of effective diffusion constants at 20ms $D_{20ms}$ calculated from time-averaged MSD (top) and ensemble-averaged MSD (bottom) in Hela cells. Mean values and standard deviations are indicated at the top-left corner of each panel. **e**. Ergodicity breaking parameters (EB) were plotted at various time delays for GEM trajectories in both the cytoplasm and nucleus of Hela cells.

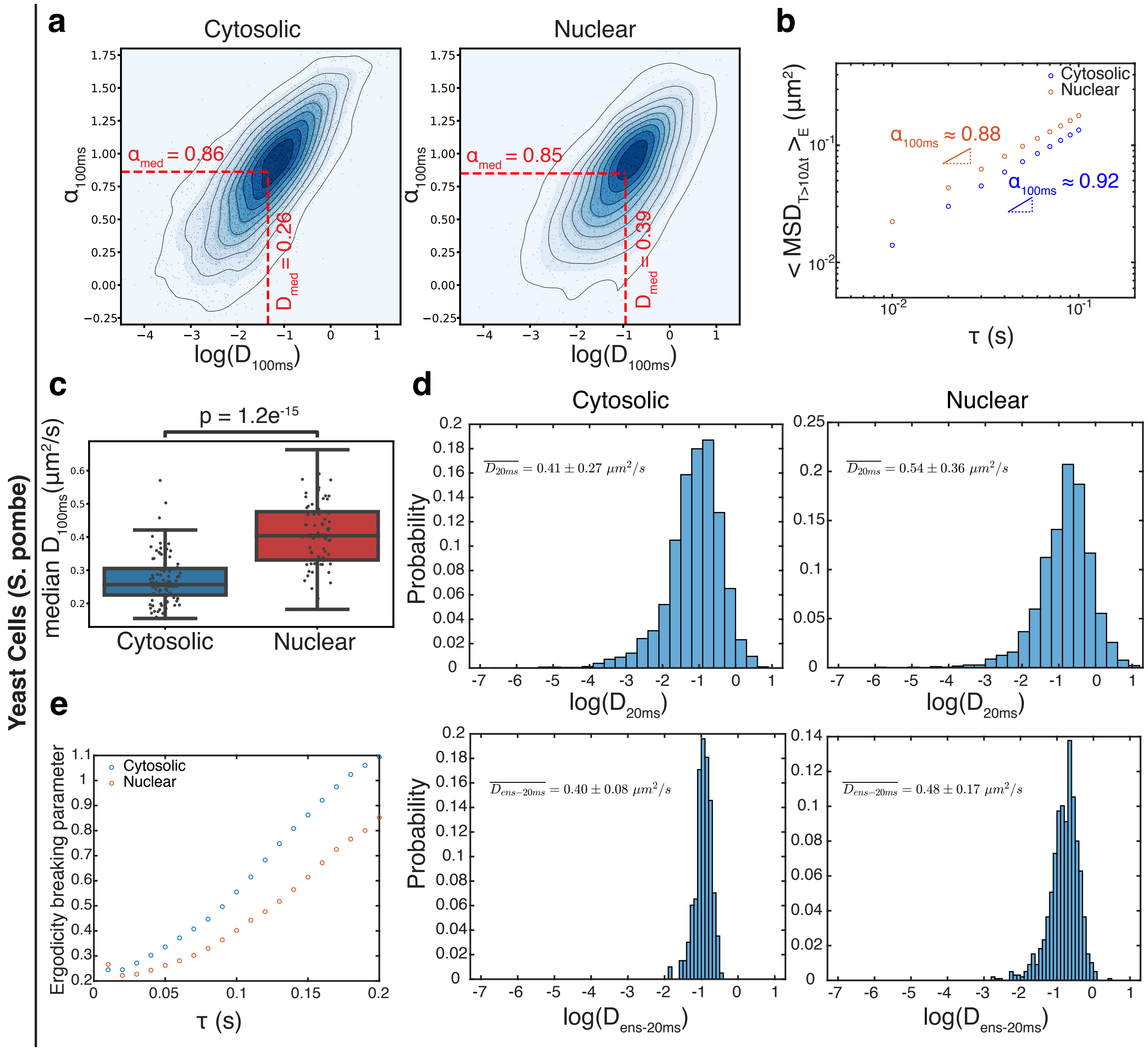


**Supplementary Figure 9: Quantification of the rheological properties of the cytosol and nucleoplasm of *Schizosaccharomyces pombe* cells. a.** Density plot of $\alpha_{100ms}$ - versus $\log(D_{100ms})$ for individual GEM trajectories in the cytoplasm (left) and nucleus (right) of *S. pombe* yeast cells. **b.** Ensemble-time-averaged MSD with fitted $\alpha_{100ms}$ values for GEM trajectories in *S. pombe* cells with larger than 10 time points. In **a-b**, n =4212 (cytosolic) and n = 1843 (nuclear) trajectories **c.** Box plot of the median effective diffusion coefficients ($D_{100ms}$) of trajectories from individual *S. pombe* yeast cells; n = 94 (cytosolic) and n = 65 (nucleus). **d.** Histograms of effective diffusion coefficients at 20ms $D_{20ms}$ calculated from time-averaged MSD (top) and ensemble-averaged MSD (bottom) in *S. pombe*. Mean values and standard deviations were labeled at the top-left corner in each figure. **e.** Ergodicity breaking parameters (EB) were plotted at various time delays for GEM trajectories in both the cytoplasm and nucleus in *S. pombe*.

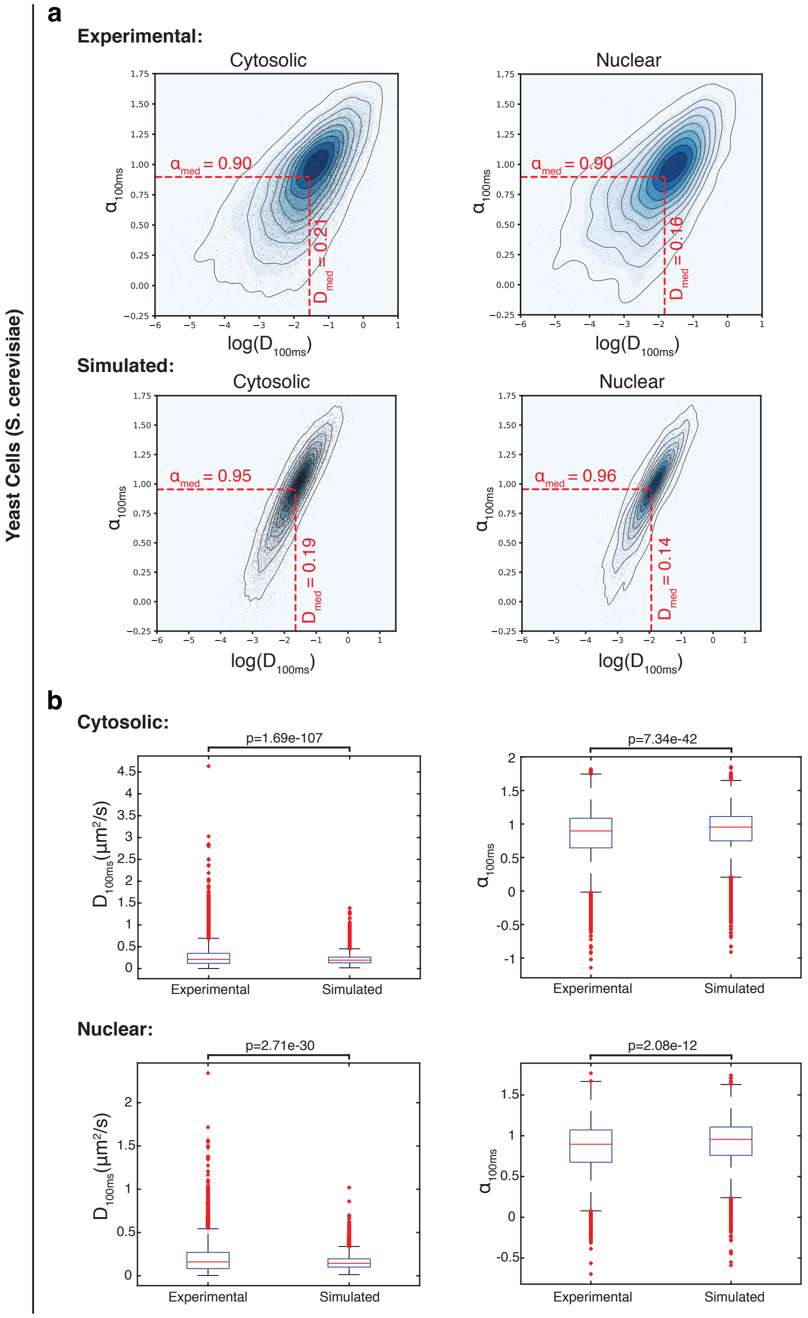


**Supplementary Figure 10: Comparison of *S. cerevisiae* experimental GEM motion to simulated Brownian motion**.

**a**. Top: Density plot of effective diffusion coefficient $log(D_{100ms})$, versus anomalous exponent $\alpha_{100ms}$, for individual GEM trajectories having more than 10 time points both in cytosol (left) and nucleus (right) of S. cerevisiae, with their median values highlighted by red dashed lines (same data as Fig. **4a**); Bottom: Brownian motion trajectories were simulated both in the cytosol (left) and nucleoplasm (right) with diffusion coefficients fixed to the experimental values. New median values were indicated as red dashed lines. Both the numbers of simulated trajectories as well as trajectory lengths are the same as in the experimental data. For both experimental and simulated data, there are n=10,706 (cytosolic) and n=2,969 (nuclear) trajectories. **b**. Box plot comparing the calculated effective diffusion coefficients ($D_{100ms}$) and anomalous exponents ($\alpha_{100ms}$) between experimental data and simulated data in the cytosol (top) and nucleoplasm (bottom). P-values were derived from Student's t-test to assess statistical differences between experimental and simulation data.

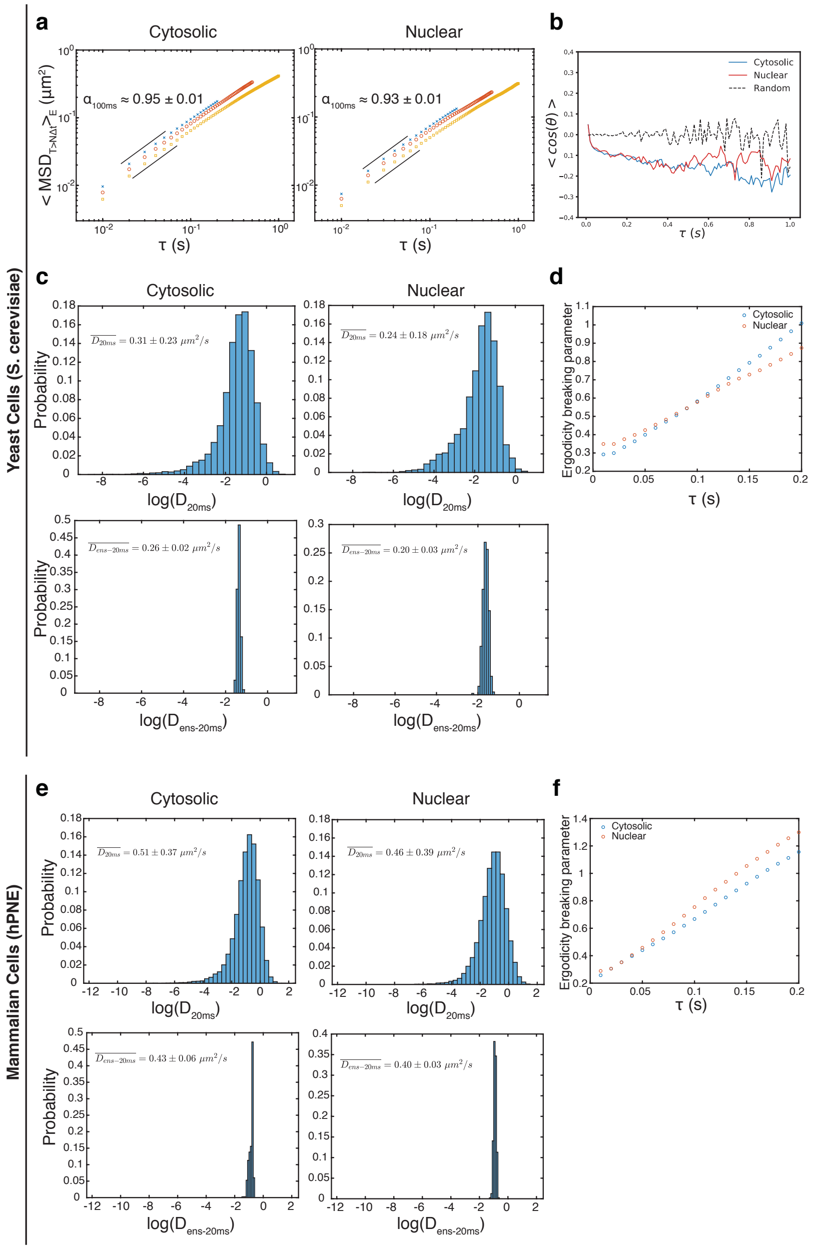


**Supplementary Figure 11: Quantification of nucGEM motion reveals distinct mesoscale rheological properties of the nucleoplasm and cytosol in both *S. cerevisiae* and mammalian pancreatic cells (hPNE). a.** Ensemble- and time-averaged mean squared displacements (MSD) versus time-step ($\tau$) with track length cutoffs of 20 (blue, $n_{cytosolic}$ = 5,355, $n_{nuclear}$ =1,589), 50 (red, $n_{cytosolic}$ = 1,848, $n_{nuclear}$ = 613), and 100 (yellow, $n_{cytosolic}$ = 622, $n_{nuclear}$ = 229) for GEM trajectories in both the cytosol and nucleoplasm of *S. cerevisiae*. Mean and standard deviation of fitted $\alpha_{100ms}$ values are shown. **b.** Angle correlation (mean cosine) functions for GEM trajectories in the nucleus (red) and cytosol (blue) of *S. cerevisiae* as a function of time-scale (τ) as well as simulated Brownian motion with uniformly distributed random angle series (dashed black). The number of angles simulated for the random angle series was matched to the number extracted at each $\tau$ for the nuclear trajectories. **c, e.** Histograms of effective diffusion coefficients at 20ms ($D_{20ms}$) calculated from time-averaged MSD (top) and ensemble-averaged MSD (bottom) in *S. cerevisiae* (**c**) and hPNE cells (**e**). Mean values and standard deviations were labeled at the top-left corner in each figure. **d, f.** Ergodicity breaking parameters (EB) were plotted at each time delay $\tau$ for GEM trajectories in both cytoplasm and nucleus in *S. cerevisiae* (**d**) and hPNE cells (**f**).

**Supplementary Video 1: *S. cerevisiae* cells expressing cytGEMs and nucGEMs.** 100 Hz, 4 second video-micrographs (HILO imaging) showing cytoGEMs (left, cyan), and nucGEMs (right, cyan) in cells with the nuclear envelope marked by Nup49 (magenta) in *Saccharomyces cerevisiae*. Each frame represents 10 msec, scale bar indicates 5 µm. Playback is in real time.

**Supplementary Video 2**: **Human pancreatic (hPNE) cells expressing cytGEMs and nucGEMs.** 100 Hz, 4 second video-micrographs (spinning disk confocal microscopy) of cytoGEMs (top, cyan), and nucGEMs (bottom, cyan) with SiR-DNA dye to indicate the position of the nucleus in human hPNE cells. Each frame represents 10 ms, scale bar indicates 10 µm. The playback is in real time.

**Supplementary Video 3: HeLa cell expressing nucGEM undergoing mitosis:** A concatenation of nine 2-second, 100 Hz video-micrographs from the same dividing HeLa cell expressing nucGEMs (cyan) and stained with SiR-DNA (magenta). Each video was acquired 15 minutes apart. The scale bar represents 10 µm and each frame rate is 10 ms.
